# Two structurally mobile regions control the conformation and function of metamorphic meiotic HORMAD proteins

**DOI:** 10.1101/2024.08.05.606648

**Authors:** Consuelo Barroso, Josh P Prince, Punam Rattu, Daimona Kundé, Nuria Ferrandiz, Syma Khalid, Enrique Martinez-Perez

## Abstract

Metamorphic HORMA domain proteins (HORMADs) nucleate protein complex formation by refolding their mobile safety belt region to bind short motifs on interactors. Meiotic HORMADs (mHORMADs) bind proteinaceous axial elements to orchestrate complex chromosomal events that underpin fertility, including pairing and recombination between homologous chromosomes. However, the mechanisms supporting the diverse roles of mHORMADs remain unclear. Here, we show that mHORMADs have a second structurally mobile region, the β5-αC loop, which controls mHORMAD conformation and function. Molecular dynamics and in vivo approaches show that functional specialisation of *C. elegans* paralogs HTP-1 and HTP-2 depends on the interplay between their β5-αC loop and safety belt. The β5-αC loop can interact with the same HORMA core surface as the safety belt, forming a “loop engaged” conformation. The β5-αC loop HORMA core interaction is essential for axis loading of HTP-1 and its paralog HTP-3, and is also present in yeast, plant, and mammalian mHORMADs, suggesting that it represents a conserved functional feature of mHORMADs. Our study reveals that mHORMADs have expanded the bimodal folding landscape first identified in Mad2, paving the way to elucidate how non-canonical HORMAD conformations control meiotic chromosome function to ensure fertility.

## Introduction

Accurate formation of haploid gametes from diploid germ cells during meiosis is essential for the fertility of sexually-reproducing organisms. Key to this process is the formation of inter-homologue crossover events during the prolonged prophase of the first meiotic division ^1^. Crossovers, together with sister chromatid cohesion provided by the cohesin complex, provide the basis of chiasmata, physical attachments between homologous chromosomes that ensure their correct orientation on the first meiotic spindle ^2^. Defects in the processes that promote chiasma formation result in sterility and the formation of aneuploid gametes, a cause of birth defects and miscarriages in humans.

The formation of chiasmata requires deliberate induction of DNA double strand breaks (DSBs) to initiate meiotic recombination, pairing and synapsis between homologous chromosomes, and repair of DSBs using the homologue as a repair template. These events in turn depend on the assembly of proteinaceous axial elements containing cohesin and a group of conserved proteins characterised by the presence of a HORMA domain (HORMADs) at the onset of meiotic prophase ^3^. Meiotic HORMADs (mHORMADs) are essential for key chromosomal transactions of meiosis, including: DSB formation, pairing and synapsis of homologous chromosomes, repair of DSBs, and quality control mechanisms that monitor pairing and recombination intermediates to control meiotic progression ^4^. Our mechanistic understanding of how mHORMADs promote this series of meiotic events is limited, but, similar to other HORMAD proteins, mHORMADs are thought to act as platforms for the assembly of protein complexes, thus the timing of their activity must be tightly regulated. Unicellular eukaryotes express a single mHORMAD, exemplified by Yeast Hop1 ^5^, but some multicellular organisms have undergone expansion of this protein family, including mammals (HORMAD1 and HORMAD2) ^6–8^ and *C. elegans* (HTP-3, HTP-1, HTP-2, and HIM-3) ^9–12^. In vivo mutant analysis provides evidence for functional specialisation between mHORMAD paralogs in these organisms ^13–17^, but a mechanistic understanding of how these mHORMAD paralogs displaying highly identical amino acid sequences and protein structure have acquired functional specialisation is currently lacking.

The HORMA domain consists of a short and flexible N-terminus, a stable core of three α helices (α A-C) and a three-stranded β-sheet (β4-6), plus a flexible C-terminal domain called the safety belt. The two β sheets situated at the edges of the core (β6 and β5) serve as interaction surfaces for the flexible safety belt, giving HORMADs the ability to acquire at least two different folding configurations that determine the activity status of the protein ^18^. This two-state behaviour is best understood for the spindle assembly checkpoint protein Mad2. Mad2 exists in an inactive and partner-free open conformation in which the safety belt interacts with β6 on the side of the core, and a partner-bound closed conformation where the safety belt moves across the HORMA core to interact with β5 and to wrap around a 6-10 amino acid closure motif (CM) on an interacting partner, thereby forming active Mad2-containing protein complexes ^19,20^. This two-state behaviour is also present in Rev7 ^21^, a HORMAD component of different DNA repair complexes ^22^. Similar to Mad2 and Rev7, mHORMADs also acquire a closed conformation bound to CMs and although Hop1 carries a chromatin-binding domain on its C-terminus ^23,24^, binding a CM on axis-bound interactors is the main mechanism by which mHORMADs are recruited to axial elements ^25,26^. However, in contrast to Mad2 and Rev7, a stable open conformation has not been identified in mHORMADs ^18^, which adopt a partially-unfolded unbuckled conformation where the safety belt is disengaged from the core of the HORMA domain ^26^. In addition, the flexible loop connecting β5 to αC is typically longer in mHORMADs than in Mad2 ^25^ and its deletion has opposing conformational effects in Mad2 (stabilisation of an open conformation^19^) and yeast Hop1 (stabilisation of a closed conformation ^26^). Therefore, although the two-state behaviour of Mad2 is thought of as a paradigm for HORMADs, the conformation dynamics of mHORMADs and how this correlates with their different functions remains poorly understood.

In contrast to Mad2, which consists exclusively of the HORMA domain, mHORMADs have additional N- and C-terminal domains flanking their HORMA domain that act as recruiting platforms for proteins controlling multiple aspects of meiotic chromosome metabolism ^25,27–29^. Current models to explain mHORMADs’ mode of action suggest that the main role of the HORMA domain is targeting of the protein to axial elements by binding CMs on axis-bound proteins, while the extended C-terminus serves as a platform to recruit proteins that drive pairing, recombination, and checkpoint control. Then, removal of mHORMADs from the axis, a process that in most organisms is mediated by the ATPase remodeller protein Pch2 (Trip13) ^30,31^, terminates activities driven by mHORMADs’ interactors. According to this model, the HORMA domain plays a relatively passive role once mHORMADs are bound to axial elements. However, recent reports show that the HORMA domain itself serves as an interaction surface to recruit recombination proteins to axial elements ^32,33^. Thus, the HORMA domain of mHORMADs can support multiple functions simultaneously, suggesting that its conformational landscape may be more complex than previously thought. Moreover, differences in the behaviour of the HORMA domain of highly identical mHORMAD paralogs could potentially underpin their functional specialisation.

Here, we combine functional in vivo studies with molecular dynamics modelling and protein folding prediction to investigate the functional specialisation of *C. elegans* HORMAD paralogs HTP-1 and HTP-2, which display 82% identity at the amino acid level. First, we demonstrate that major functional differences between these proteins are due to 7 amino acid changes in the safety belt region. Second, we identify the interplay between the safety belt and the extended β5-αC loop as key for controlling mHORMAD conformation and demonstrate that the β5-αC loop is essential for HTP-1 function in vivo. Third, structural predictions and in vivo analysis of *C. elegans* HTP-3 reveal that the β5-αC loop can interact with the HORMA core, forming a novel “loop engaged conformation” that is required for HTP-3 loading to chromosomes. Finally, structural prediction of mHORMADs in other organisms, including plants and mammals, suggest that the ability of the extended β5-αC loop to control the conformation of mHORMADs is a conserved feature of these proteins. We propose a model in which the position of the β5-αC loop determines different mHORMAD conformations that are essential for the successful execution of the meiotic programme and therefore for fertility.

## Results

### HTP-2 is unable to functionally replace HTP-1

HTP-1 and HTP-2 are 82% percent identical at the amino acid level and the crystal structures of their HORMA domains show that they appear nearly identical ^25^. Despite this, previous analysis of single and double mutant worms carrying the *htp-1*(*gk174*) and *htp-2*(*tm2543*) alleles suggested that HTP-2 is unable to functionally replace HTP-1. While meiosis appears largely unaffected in *htp-2*(*tm2543*) mutants, *htp-1*(*gk174*) mutants display defects in homologue pairing, SC assembly, recombination, and checkpoint control of meiotic progression ^10,15^. Analysis of *htp-1*(*gk174*) *htp-2*(*tm2543*) double mutants suggests that HTP-2 supports homology-independent SC assembly and together with HTP-1 controls the spatial release of cohesin during the first meiotic division ^13,27^. In addition, immunostaining with antibodies recognising both HTP-1 and HTP-2 in wild-type (WT) controls, *htp-1(gk174)* mutants, and *htp-2(tm2543)* mutants indicated that HTP-1 is more abundant than HTP-2 on pachytene chromosomes ^13^, suggesting that differences in chromosome loading may explain the inability of HTP-2 to functionally replace HTP-1. To clarify the contribution of HTP-1 and HTP-2 to crossover formation and whether this is determined by the relative amounts of these proteins on meiotic chromosomes, we used CRISPR to: First, create *htp-1* and *htp-2* alleles deleting the whole coding region of both genes (referred to as *htp-1Δ* and *htp-2Δ* respectively). Second, create *htp-1::FLAG* and *htp-2::FLAG* alleles carrying a FLAG tag before the STOP codon. Third, substitute the *htp-1* gene (sequence between start codon and STOP codon) with the corresponding sequence from the *htp-2* gene, resulting in worms that carry two copies of *htp-2* (referred to as *twice htp-2*) in which the additional copy of *htp-2* is expressed under the endogenous *htp-1* promoter and 3’ UTR.

The *C. elegans* genome contains six pairs of homologous chromosomes, which are observed as six DAPI-stained bodies in diakinesis oocytes of WT worms due to the formation of crossover events between all pairs of homologous chromosomes (Figure 1a). In contrast, failure in crossover formation manifests in the presence of seven to twelve DAPI-stained bodies in diakinesis oocytes depending on the number of homologue pairs failing to undergo crossover formation. Oocytes of WT controls and *htp-2Δ* mutants displayed six DAPI-stained bodies, confirming that HTP-2 is not required for crossover formation (Figure 1a). In contrast, oocytes of *htp-1(gk174)*, *htp-1Δ*, and *twice htp-2* mutants displayed an average of 10.9, 11.1, and 11.2 DAPI-stained bodies respectively, indicating a severe failure in crossover formation (Figure 1a). We further corroborated this result by measuring embryonic lethality, which arises as a consequence of defective crossover formation between autosomes, and the incidence of male progeny, which provides a readout of crossover failure between the X chromosomes during meiosis. While WT controls and *htp-2Δ* mutants displayed very low levels of embryonic lethality and male progeny, *htp-1(gk174)*, *htp-1Δ*, and *twice htp-2* mutants displayed over 95% embryonic lethality and high levels of male progeny (Figure 1b). Thus, HTP-2 alone, even when simultaneously expressed from its endogenous locus and the *htp-1* locus, is not capable of supporting crossover formation. We also confirmed that, as reported for the *htp-1*(*gk174*) *htp-2*(*tm2543*) double mutant ^13^, *htp-1Δ htp-2Δ* double mutants displayed reduced and delayed SC assembly compared with *htp-1Δ* single mutants (Figure S1a), consistent with HTP-2 supporting homology-independent SC assembly.

**Figure 1.**
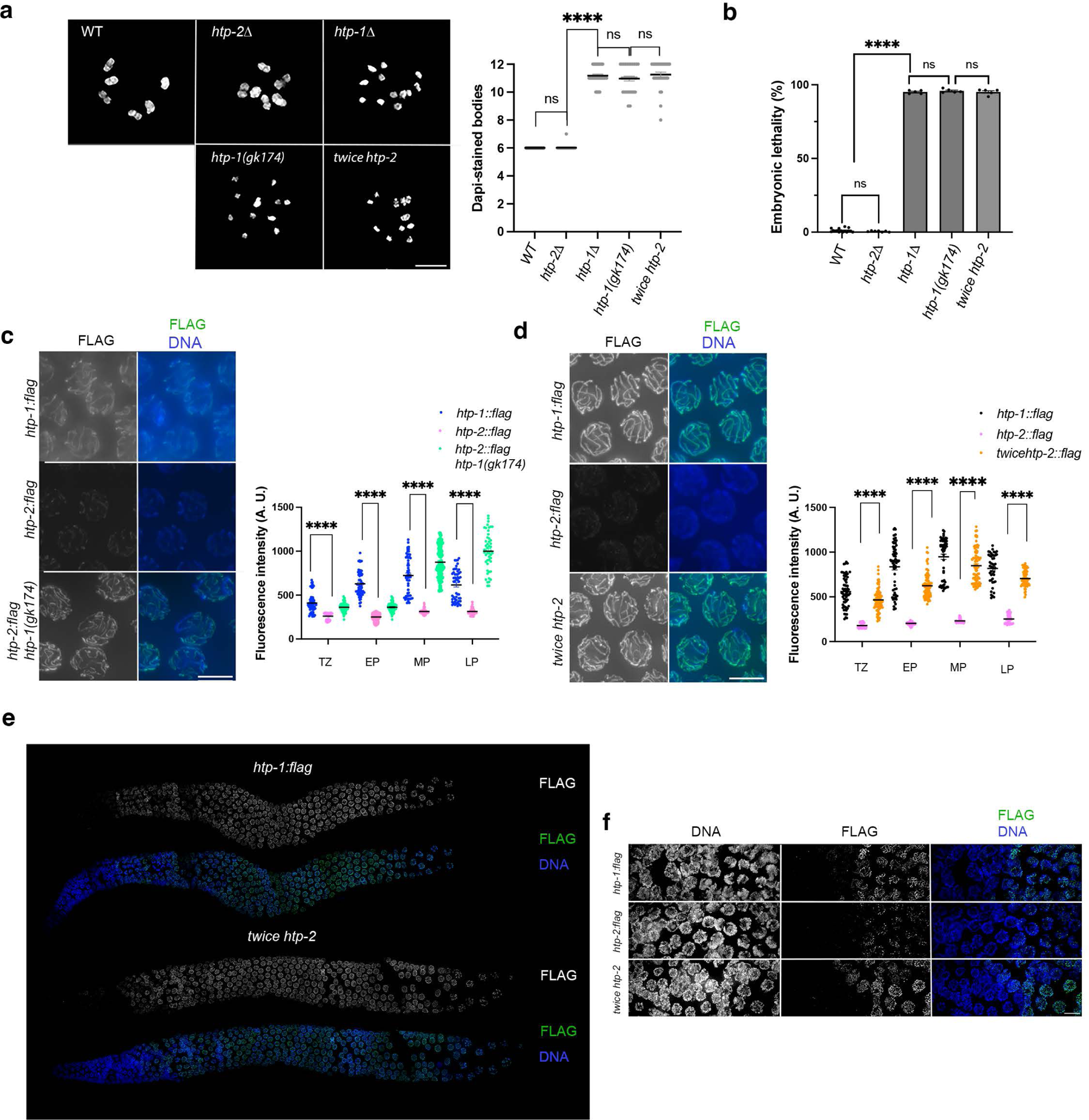
HTP-1 and HTP-2 are functionally distinct. **a)** Projections of diakinesis oocytes from indicated genotypes stained with DAPI. 6 DAPI-stained bodies indicate normal crossover formation, while complete absence of crossover results in 12 DAPI-stained bodies. Graph shows quantification of number of DAPI-stained bodies per genotype (n= 30 (WT); n= 65 (*htp-2Δ*); n= 35 (*htp-1Δ*); n= 30 (*htp-1(gk174)*); n= 31 (*twice htp-2*), error bars indicate mean with 95% CI, p values were calculated using a two-tailed Mann-Whitney U test. Note large crossover failure in *htp-1Δ*, *htp-1(gk174),* and *twice htp-2* mutants. **b)** Quantification of embryonic lethality in strains of indicated genotypes. Number of worms and embryos analysed per genotype: WT= 15, 3421; *htp-2Δ*= 7, 1842; *htp-1Δ*= 5, 878; *htp-1(gk174)*= 5, 727; *twice htp-2*= 5, 1165, error bars indicate mean with 95% CI, p values were calculated using a two-tailed Mann-Whitney U test. **c-d)** Non-deconvolved projections of pachytene nuclei from indicated genotypes stained with anti-FLAG antibodies and DAPI. Images were acquired and adjusted with the same settings for all genotypes. Graphs show intensity of anti-FLAG staining in nuclei of the indicated germline regions (transition zone (TZ), early pachytene (EP); mid pachytene (MP), and late pachytene (LP) and genotypes). Between 49 and 83 nuclei from three to four different germlines were analysed per stage and genotype, error bars indicate mean with 95% CI, p values were calculated using a two-tailed Mann-Whitney U test. **e)** Deconvolved projections of whole germlines of indicated genotypes stained with anti-FLAG antibodies and DAPI. Note similar pattern of anti-FLAG staining in *htp-1::FLAG* and *twice htp-2::FLAG* (in this case FLAG signal corresponds to HTP-2 expressed from the *htp-1* locus) germlines. **f)** Deconvolved images of transition zone nuclei from indicated genotypes showing that FLAG associates with similar intensity and timing in *htp-1::FLAG* and *twice htp-2::FLAG* germlines. Scale bar =5 µm in all panels.

Next, we used the FLAG-tagged versions of *htp-1* and *htp-2* in immunostaining experiments to determine the relative abundance of HTP-1 and HTP-2 and to compare the loading pattern of both proteins through meiotic prophase. Worms homozygous for *htp-1::FLAG* or *htp-2::FLAG* in otherwise WT genetic backgrounds displayed normal levels of crossovers and embryonic viability, confirming that FLAG tagging did not affect protein function (Figure S1b-c). Comparing anti-FLAG staining intensity in germlines of *htp-1::FLAG* and *htp-2::FLAG* homozygous worms demonstrated that during WT meiosis the amount of HTP-1 associated with meiotic chromosomes was significantly higher than that of HTP-2 at all stages of meiotic prophase (Figure 1c). To determine if the presence of HTP-1 affects the loading of HTP-2, we also measured the intensity of HTP-2::FLAG staining in germlines lacking HTP-1. Removal of HTP-1 caused a clear increase in the amount of HTP-2 associated with meiotic chromosomes, especially in nuclei at later stages of meiotic prophase (Figure 1c). Although HTP-2::FLAG levels were also increased in transition zone and early pachytene nuclei of *htp-1* mutants, the intensity of the FLAG signal did not reach levels observed for that of HTP-1::FLAG (Figure 1c). In contrast to the large increase in HTP-2 loading seen in *htp-1* mutants, removing HTP-2 only induced modest changes in HTP-1 levels (Figure S1d). Thus, HTP-2 loading is limited by HTP-1 but not vice versa.

Given that HTP-1 plays key roles in ensuring correct homologue pairing and synapsis during early meiotic prophase ^14,15^, we wondered whether increasing the levels of chromosome-bound HTP-2 during early prophase would be sufficient to functionally replace HTP-1. Thus, we also imaged germlines from *twice htp-2* mutants in which we added a FLAG tag to the *htp-2* copy expressed from the *htp-1* locus, while the endogenous *htp-2* locus remained untagged. These germlines displayed a striking increase in the intensity of HTP-2::FLAG compared to WT *htp-2::FLAG* worms (Figure 1d). In fact, the staining intensity and loading pattern of HTP-2::FLAG in germlines of *twice htp-2* were very similar to the intensity levels and loading pattern of HTP-1::FLAG observed in WT *htp-1::FLAG* germlines, including in transition zone nuclei (Figures 1e-f). Crucially, however, *twice htp-2* mutants were not competent in crossover formation (Figure 1a). Therefore, increasing the amount of HTP-2 that associates with chromosomes from the onset of meiosis is not sufficient to functionally replace HTP-1, demonstrating that HTP-1 and HTP-2 are functionally divergent. In addition, our observations also demonstrate that HTP-1 competes with HTP-2 for loading onto axial elements.

### Functional differences between HTP-1 and HTP-2 map to the C-terminal region of the HORMA domain

Having found that HTP-2 is unable to provide HTP-1 functions needed for crossover formation, we reasoned that functional differences between these proteins must be conferred by differences in specific amino acids. In order to map the regions containing these amino acids, we decided to create a series of chimeric HTP-1 proteins in which different domains were replaced with the corresponding HTP-2 sequence. These chimeric proteins were expressed from a single-copy transgene containing the *htp-1* promoter and 3’ UTR and crossed into a *htp-1(gk174)* mutant background to determine whether the chimeric protein supports crossover formation. Meiotic HORMA-domain proteins, including HTP-1 and HTP-2, consist of three broad domains: A short N-terminal domain, the HORMA domain, and a C-terminal domain typically containing a closure motif (Figure 2a). As the 60 amino acid differences (out of 352) between HTP-1 and HTP-2 are distributed relatively evenly between the different protein domains (Figure 2b), we initially created chimeric proteins swapping the N-terminal (HTP-1^HTP-2 N-term^), HORMA (HTP-1^HTP-2 HORMA^), or C-terminal (HTP-1^HTP-2 C-term^) domains (Figure 2a). Next, we assessed the functionality of these proteins by measuring embryonic lethality and number of DAPI-stained bodies in diakinesis oocytes of worms homozygous for the different transgenes and the *htp-1(gk174)* allele. A transgene expressing a WT version of HTP-1 was included as a positive control. *htp-1^htp-2 N-term^* worms displayed levels of embryonic lethality and numbers of DAPI-stained bodies similar to WT controls (Figures 2c-d), evidencing that the N-terminus of HTP-1/2 are functionally interchangeable. In contrast, worms expressing HTP-1^HTP-2 HORMA^ displayed 88% embryonic lethality and an average of 9.4 DAPI-stained bodies (Figures 2c-d), demonstrating that the HORMA domain of HTP-2 does not support HTP-1 function. Finally, *htp-1^htp-2 C-term^* worms displayed 36 % embryonic lethality and a slight increase in the number of DAPI-stained bodies (average of 6.6 versus 6 in controls) (Figures 2c-d), suggesting the presence of mild meiotic defects. These experiments suggest that amino acid substitutions within the HORMA domain may be responsible for functional differences between HTP-1 and HTP-2.

**Figure 2.**
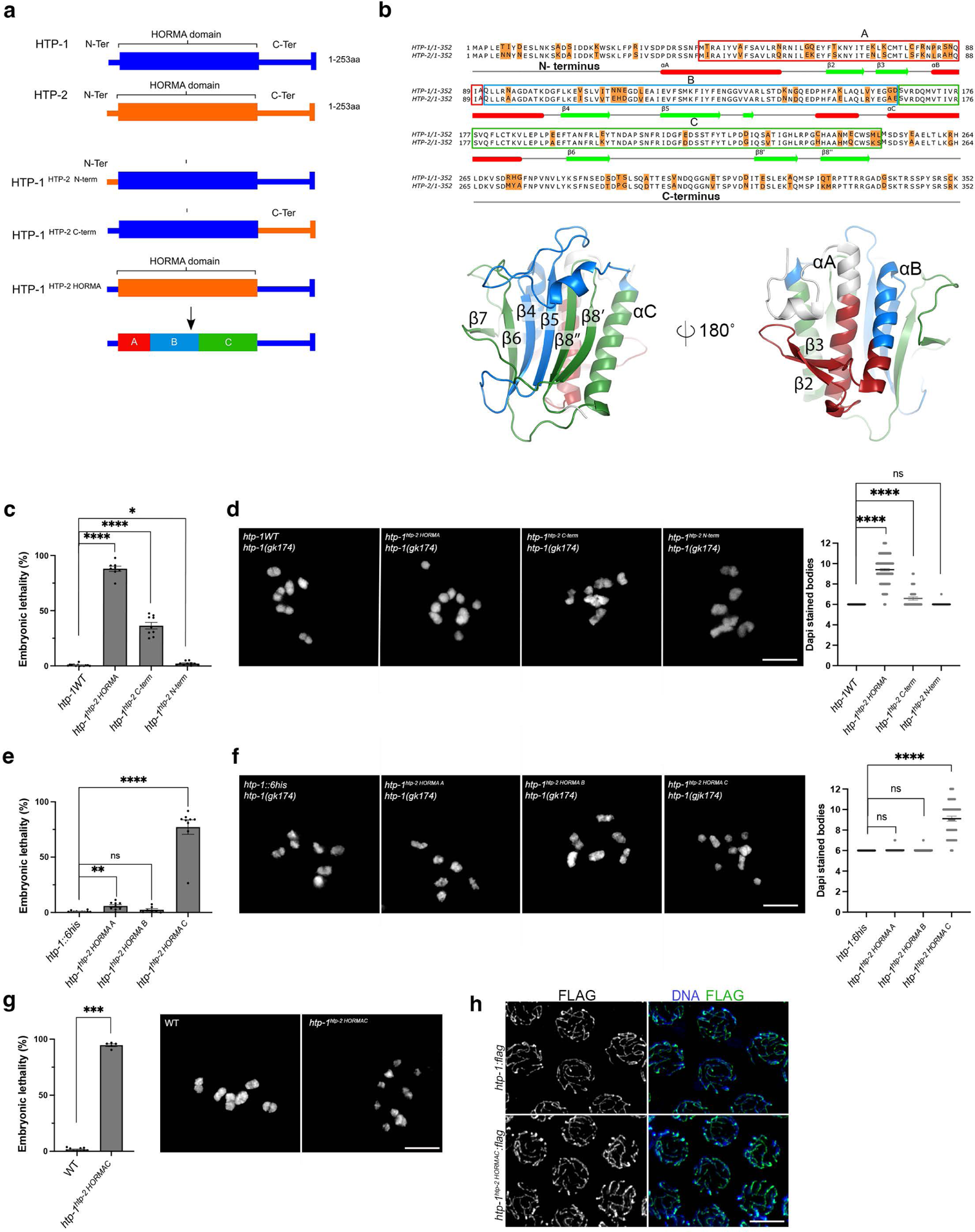
Mapping functional differences between HTP-1 and HTP-2. **a)** Cartoon depicting the three broad domains (N-terminus, HORMA domain, and C-terminus) present in HTP-1 and HTP-2 and chimeric proteins in which HTP-1 domains were replaced with their HTP-2 counterpart. Bottom cartoon shows the three subdomains (A,B, C) in which the HORMA domain was divided. **b)** Alignment of HTP-1 and HTP-2 indicating amino acid substitutions (highlighted in orange), secondary structure features of both proteins (CM indicates closure motif), and the amino acids included in the A (red outline), B (blue outline), and C (green outline) HORMA subdomains. Bottom panel shows structural features of HTP-1/2 HORMA domain indicating the HORMA subdomains in red, blue, and green. **c)** Quantification of embryonic lethality in strains of indicated genotypes. Number of worms and embryos analysed per genotype: *htp-1^WT^ htp-1(gk174)*= 13, 3685; *htp-1^htp-2 HORMA^ htp-1(gk174)*= 8, 1864; *htp-1^htp-2C-term^ htp-1(gk174)*= 9, 1888; *htp-1^htp-2 N-term^ htp-1(gk174)*= 10, 2835, error bars indicate mean with 95% CI, p values were calculated using a two-tailed Mann-Whitney U test. **d)** Quantification of number of DAPI-stained bodies in diakinesis oocytes of worms of indicated genotypes. Number of oocytes analysed *htp-1^WT^ htp-1(gk174)*= 39; *htp-1^htp-2 HORMA^ htp-1(gk174)*= 135; *htp-1^htp-2C-term^ htp-1(gk174)*= 34; *htp-1^htp-2 N-term^ htp-1(gk174)*= 60, error bars indicate mean with 95% CI, p values were calculated using a two-tailed Mann-Whitney U test. **e)** Quantification of embryonic lethality in strains of indicated genotypes. Number of worms and embryos analysed per genotype: *htp-1^WT^ htp-1(gk174)*= 10, 2637; *htp-1^htp-2 HORMA A^ htp-1(gk174)*= 9, 2415; *htp-1^htp-2 HORMA B^ htp-1(gk174)*= 7, 1578; *htp-1^htp-2 HORMAC^ htp-1(gk174)*= 9, 1590; error bars indicate mean with 95% CI, p values were calculated using a two-tailed Mann-Whitney U test. Note high embryonic lethality in *htp-1^htp-2 HORMAC^ htp-1(gk174).* **f)** Quantification of number of DAPI-stained bodies in diakinesis oocytes of worms of indicated genotypes. Number of oocytes analysed *htp-1^WT^ htp-1(gk174)*= 36; *htp-1^htp-2 HORMA A^ htp-1(gk174)*= 42; *htp-1^htp-2 HORMA B^ htp-1(gk174)*= 59; *htp-1^htp-2 HORMAC^ htp-1(gk174)*= 46; error bars indicate mean with 95% CI, p values were calculated using a two-tailed Mann-Whitney U test. Note the increased number of DAPI-stained bodies present in *htp-1^htp-2 HORMAC^ htp-1(gk174)* oocytes compared to WT controls. **g)** *htp-1^htp-2 HORMAC^* mutants created by CRISPR display elevated embryonic lethality (left panel) and increased numbers of DAPI-stained bodies in diakinesis oocytes (right panel). Numbers of worms and embryos analysed per genotype WT= 10, 2496; *htp-1^htp-2 HORMAC^*= 5, 819. Number of oocytes analysed WT= 30; *htp-1^htp-2 HORMAC^*= 35. Error bars indicate mean with 95% CI, p values were calculated using a two-tailed Mann-Whitney U test. **h)** Deconvolved projections of pachytene nuclei of indicated genotypes stained with anti-FLAG antibodies and DAPI. Note similar anti-FLAG staining pattern in WT controls and *htp-1^htp-2 HORMAC^* mutant nuclei. Scale bar =5 µm in all panels.

Next, we divided the HORMA domain into 3 regions (HORMA A, B, and C) containing an even distribution of amino acid diferences (13, 13, and 10 respectively) (Figures 2a-b) and created worms expressing the corresponding chimeric HTP-1/HTP-2 proteins from a single copy transgene in a *htp-1(gk174)* background as explained above. *htp-1^htp-2 HORMA A^*, *htp-1^htp-2 HORMA B^,* and *htp-1^WT^* worms displayed low levels of embryonic lethality (Figure 2e) and were competent in crossover formation as indicated by the number of DAPI-stained bodies in diakinesis oocytes (Figure 2f). In contrast, *htp-1^htp-2 HORMA C^* worms displayed high levels of embryonic lethality (Figure 2e) and an average of 9.1 DAPI-stained bodies in diakinesis oocytes (Figure 2f), demonstrating a severe crossover defect. Therefore, residues between E191 and L250, which expand the ß6 to ß8’’ regions of HTP-1’s HORMA domain (Figure 2b) are critical for the ability of HTP-1 to ensure crossover formation. To corroborate this finding, we used CRISPR to introduce the 10 amino acid substitutions contained in the *htp-1^htp-2 HORMA C^* transgene into the endogenous *htp-1* locus. *htp-1^htp-2 HORMA C^* (CRISPR) mutants displayed 95% embryonic lethality and displayed univalents in diakinesis oocytes (Figure 2g), replicating our observations of worms expressing the *htp-1^htp-2 HORMA C^* transgene in a *htp-1(gk174)* background. We also used CRISPR to introduce a FLAG tag before the STOP codon of HTP-1^HTP-2 HORMA C^ in order to monitor its loading to meiotic chromosomes. Anti-FLAG staining confirmed that HTP-1^HTP-2 HORMA C^::FLAG loaded to chromosomes throughout meiotic prophase (Figure 2h). Therefore, the 10 amino acid substitutions present in HTP-1^HTP-2 HORMA C^ preclude HTP-1 from ensuring normal crossover formation despite substantial loading to meiotic chromosomes.

### HTP-1^HTP-2 HORMA C^ is deficient in promoting pairing, synapsis, and checkpoint regulation of meiotic progression

HTP-1 ensures crossover formation by participating in multiple meiotic events, including initial homologue recognition, SC assembly, checkpoint regulation of meiotic progression, and formation and repair of DSBs ^14,15^. Thus, we next sought to clarify if the crossover defect observed in *htp-1^htp-2 HORMA C^* mutants was due to defects in these events. WT and *htp-1(gk174)* mutants were included as positive and negative controls respectively. We also included *twice htp-2* mutant worms in these experiments to clarify if increased levels of HTP-2 loading could improve any of the defects observed in *htp-1(gk174)* mutants, where only HTP-2 expressed from the *htp-2* locus is present. First, we assessed homologue pairing using fluorescence in situ hybridisation (FISH) to monitor the pairing of an interstitial region on chromosome V between the onset of meiotic prophase (transition zone nuclei) and late pachytene by dividing this section of the germline into six equal-size regions and counting the number of nuclei with paired signals in each zone (Figure 3a). As expected, in WT control germlines pairing levels reached nearly a 100% for most of meiotic prophase (Figure 3a). In contrast, *htp-1(gk174)*, *twice htp-2*, and *htp-1^htp-2 HORMA C^* mutants displayed severely reduced levels of pairing compared to WT controls (Figure 3a). Interestingly, pairing levels were slightly higher in *htp-1^htp-2 HORMA C^* mutants than in *htp-1(gk174)* and *twice htp-2* mutants, suggesting that HTP-1^HTP-2 HORMA C^ retains a low level of pairing-promoting activity. Next, we used antibodies against axis protein HIM-3 ^34^ and SC component SYP-1 ^35^ to monitor the progression of synapsis between transition zone and late pachytene by dividing this region of the germline into five equal-size regions. Nuclei displaying HIM-3 stretches that did not colocalise with SYP-1 staining were scored as unsynapsed, while nuclei in which all HIM-3 stretches overlapped with SYP-1 signal were scored as synapsed. In WT controls the number of nuclei displaying full synapsis increased as nuclei progress into pachytene, reaching nearly 100 % by mid pachytene before decreasing as nuclei start the process of desynapsis in late pachytene (Figure 3b). In contrast, the percentage of nuclei displaying full synapsis in *htp-1^htp-2 HORMA C^* mutants never rose above 19% and overall was very similar to the pattern observed in *twice htp-2* mutants, evidencing a clear defect in SC assembly in both mutants (Figure 3b).

**Figure 3.**
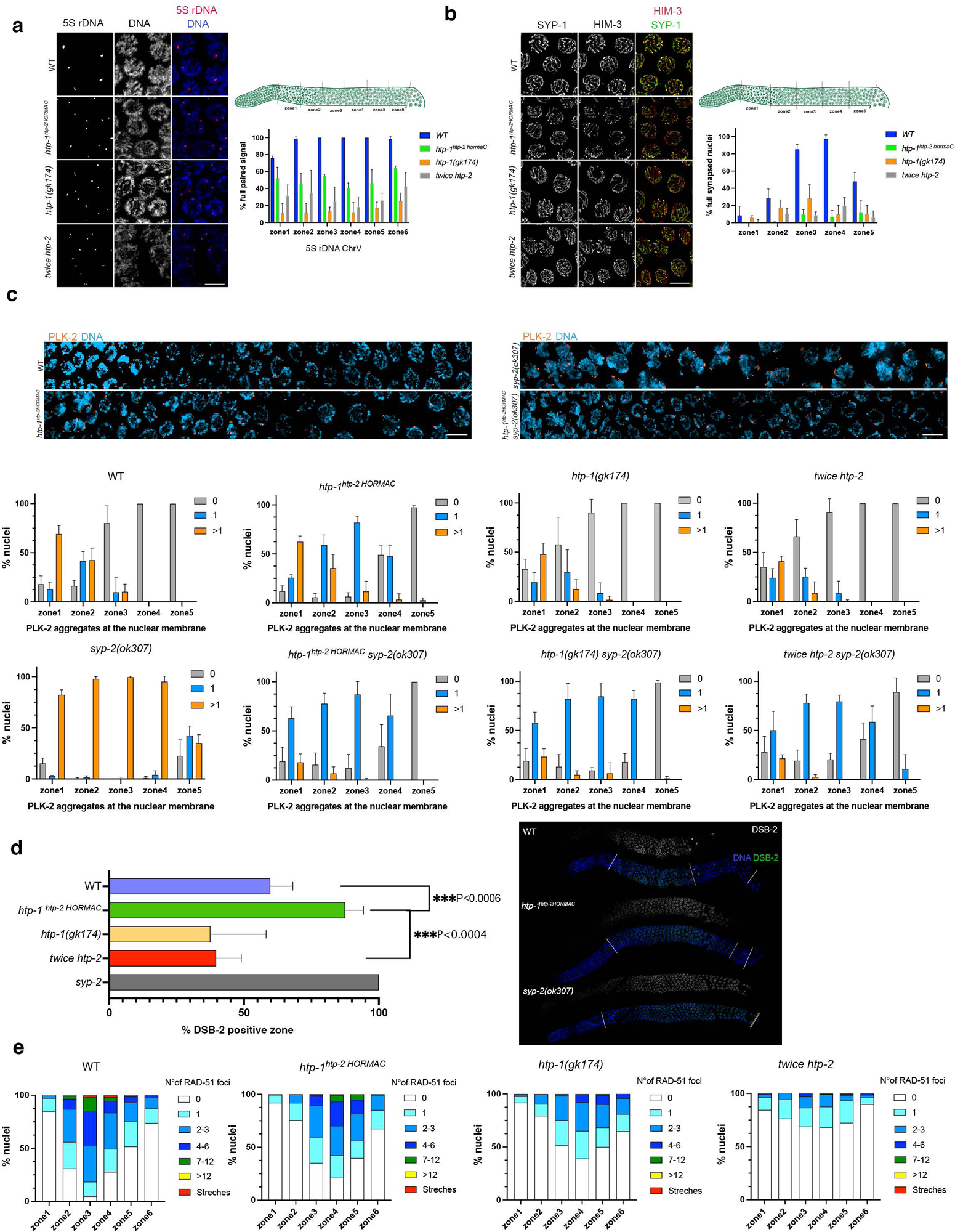
*htp-1^htp-2 HORMAC^* mutants display multiple meiotic defects. **a)** Projections of pachytene nuclei following labelling with a FISH probe to visualise the 5S rDNA locus and DAPI. Two foci per nucleus indicates a defect in homologue pairing. Graph shows quantification of the % of nuclei with paired signals (1 focus per nucleus) in 6 zones along the germline as indicated in top cartoon. Number of nuclei (3 to 4 germlines per genotype) analysed per zone= 124, 83, 72, 77, 64, 62 (WT); 107, 98, 90, 75, 81, 83 (*htp-1^htp-2 HORMAC^*); 100, 84, 70, 69, 74, 94 (*htp-1(gk174));* 99, 78, 63, 45, 49, 66 (*twice htp-2*). Note reduced pairing levels in all *htp-1* mutants. **b)** Projections of pachytene nuclei stained with anti-HIM-3 (axial element) and anti-SYP-1 (SC component) antibodies. Lines displaying only HIM-3 staining (red signal) indicate the presence of unsynapsed regions. Graph shows quantification of the percentage of nuclei displaying full synapsis (complete overlap of HIM-3 and SYP-1 signals) in five zones along the germline as indicated in the top cartoon. Number of nuclei (3 to 5 germlines per genotype) analysed per zone= 88, 108, 97, 97, 74 (WT); 201, 248, 215, 194, 140 (*htp-1^htp-2 HORMAC^*); 119, 123, 117, 89, 79 (*htp-1(gk174));* 134, 146, 125, 104, 107 (*twice htp-2*). Note SC defect in all *htp-1* mutants. **c)** Projections of nuclei in transition zone and early pachytene regions of the germline stained with anti-PLK-2 antibodies and DAPI. Nuclei with more than 1 PLK-2 aggregate indicate high CHK-2 activity, 1 PLK-2 aggregate indicates intermediate CHK-2 activity and no PLK-2 aggregate indicates no CHK-2 activity. Graphs show quantification of the % of nuclei with a given number of PLK-2 aggregates in five zones along the germline as indicated in the cartoon shown in b. Note that all *htp-1* mutants fail to accumulate nuclei with multiple PLK-2 aggregates, even in a *syp-2* mutant background. Number of nuclei (3 to 5 germlines per genotype) analysed per zone= 185, 173, 146, 129, 123 (WT); 131, 126, 109, 88, 104 (*htp-1^htp-2 HORMAC^*); 298, 265, 283, 236, 199 (*htp-1(gk174));* 162, 142, 145, 123, 123 (*twice htp-2*); 182, 180, 150, 128, 71 (*syp-2*); 158, 132, 123, 92, 69 (*htp-1^htp-2 HORMAC^ syp-2*); 127, 101, 99, 75, 71 (*htp-1(gk174) syp-2);* 162, 139, 138, 128, 128 (*twice htp-2 syp-2*). **d)** *htp-1^htp-2 HORMAC^*, but not *htp-1(gk174)* or *twice htp-2* mutants accumulate nuclei positive for DSB-2. Graph shows the percentage of vertical rows of nuclei between transition zone and late pachytene that are positive of anti-DSB-2 staining, between 3 and 6 germlines were scored per genotype, error bars indicate mean plus SD, p values were calculated using one way ANOVA between *htp-1^htp-2 HORMAC^, htp-1 (gk174)* and *twice htp-2* and an unpair t test between *htp-1^htp-2 HORMAC^* and WT. Panel on the right shows projections of full germlines stained with anti-DSB-2 antibodies and DAPI. Vertical white lines indicate region between start of transition zone and end of late pachytene, while an orange line indicates the end of the region containing DSB-2-positive nuclei. Number of germlines scored per genotype: WT= 6, *htp-1^htp-2 HORMAC^*= 4, *twice htp-2*= 5, *htp-1(gk174)*= 4 and *syp-2*= 3. **e)** Quantification of RAD-51 foci per nucleus in germlines of indicated genotypes divided into 6 zones between transition zone and late pachytene as indicated in the cartoon in a. *htp-1^htp-2 HORMAC^* mutants display higher levels of RAD-51 foci than *htp-1(gk174)* or *twice htp-2* mutants. Number of nuclei (3 germlines per genotype) analysed per zone= 183, 148, 124, 115, 89, 88 (WT); 151, 136, 122, 118, 98, 74 (*htp-1^htp-2HORMAC^*); 136, 107, 110, 92, 92, 74 (*htp-1(gk174));* 194, 161, 135, 145, 127, 79 (*twice htp-2*). Scale bar =5 µm in all panels.

HTP-1 is an essential component of quality control mechanisms that orchestrate early meiotic progression by monitoring pairing and recombination intermediates to modulate the activity of the CHK-2 kinase, which induces DSB formation and chromosome movements required for pairing and synapsis ^14,15,36–41^. One such mechanism monitors SC assembly to delay exit from early meiotic stages characterised by high levels of CHK-2 activity, which induce chromosome movements and DSB formation, until all homologue pairs achieve synapsis. A second HTP-1-dependent mechanism monitors the presence of crossover-fated recombination events between all homologue pairs to regulate exit from DSB-formation competent stages characterised by intermediate levels of CHK-2 activity ^42,43^. These checkpoints arrest nuclei at the stage of meiotic prophase when the primary defect is identified: leptotene-zygotene (high CHK-2 activity) in the case of synapsis defects and early pachytene (intermediate CHK-2 activity) in the case of impaired recombination ^37,44^. CHK-2 promotes chromosome movements during early prophase by phosphorylating multiple targets, including pairing center-binding (PCB) proteins that associate with the chromosomal end tethered to the nuclear envelope ^37,45^. This induces recruitment of PLK-2 to PCB proteins, forming dynamic PLK-2 aggregates that modify nuclear envelope components to promote homologue pairing and synapsis ^46,47^. Autosome-associated PLK-2 aggregates are dismantled once all homologue pairs achieve synapsis, but the PLK-2 aggregate associated with the paired X-chromosomes persists until crossover-fated events are formed on all homologue pairs ^41^. Thus, mutants with synapsis defects accumulate nuclei with multiple PLK-2 aggregates, while mutants deficient in crossover formation but proficient in synapsis accumulate nuclei with a single PLK-2 aggregate.

Given the defect in SC assembly that we observed in *htp-1^htp-2 HORMA C^* mutants (Figure 3b), we reasoned that if HTP-1^HTP-2 HORMA C^ is competent in monitoring synapsis to regulate CHK-2 activity, then markers of chromosome movement should persist through the pachytene region. In WT controls, nuclei with multiple PLK-2 aggregates were most abundant in zone 1 (transition zone) and 2 (early pachytene) and mostly lacking in zones 3 to 5 (Figure 3c). Similarly, nuclei with multiple PLK-2 aggregates were largely restricted to zones 1 and 2 of *htp-1^htp-2 HORMA C^*, *htp-1(gk174)*, and *twice htp-2* mutants (Figure 3c and Figure S2a), despite the synapsis defect observed in all three mutants (Figure 3b). In contrast, *syp-2* mutants, which lack an essential component of the SC ^48^ and were used as a positive control, displayed nearly a 100% of nuclei in zones 1 to 4 with multiple PLK-2 aggregates. Interestingly, *htp-1^htp-2 HORMA C^*, but not *htp-1(gk174)* or *twice htp-2*, mutants displayed an accumulation of nuclei with a single PLK-2 focus in zones 3 and 4 (Figure 3c), suggesting persistence of intermediate levels of CHK-2 activity. As the temporal extension of CHK-2 activity in response to synapsis defects depends on the severity of the synapsis defect ^37^, we wondered whether levels of SC assembly in *htp-1^htp-2 HORMA C^*, *htp-1(gk174)*, and *twice htp-2* mutants were sufficient to prevent CHK-2 activity extension. Thus, we crossed all three mutants into a *syp-2* mutant background to eliminate SC assembly. However, complete asynapsis failed to restore the accumulation of nuclei with multiple PLK-2 aggregates in *htp-1^htp-2 HORMA C^*, *htp-1(gk174)*, and *twice htp-2* mutants (Figure 3c and Figure S2a), confirming that these mutants are unable to sustain high CHK-2 activity even in the complete absence of synapsis. Instead, the three double mutants accumulated nuclei with 1 PLK-2 focus, suggesting the persistence of intermediate levels of CHK-2 activity (Figure 3c).

Our analysis of diakinesis oocytes demonstrates a severe defect in crossover formation in *htp-1^htp-2 HORMA C^* and *twice htp-2* mutants (Figures 1a and 2g). Therefore, we also investigated if these mutants were competent in delaying exit from DSB-competent stages characterised by intermediate levels of CHK-2 activity when crossover formation is impaired. To achieve this, we monitored the presence of DSB-2, a marker of DSB competence whose removal from chromosomes is coupled to the successful formation of crossover-fated events on all chromosomes ^42^. We included *htp-1(gk174)* and *syp-2* mutants as negative and positive controls for checkpoint activity respectively. *htp-1^htp-2 HORMA C^* mutants displayed a significant expansion of DSB-2-positive nuclei compared to WT controls, while *twice htp-2* mutants showed levels similar to *htp-1(gk174)* (Figure 3d). Thus, the recombination checkpoint, which delays exit from prophase stages with intermediate levels of CHK-2 activity when crossover formation is impaired, appears to be functional in *htp-1^htp-2 HORMA C^* mutants but not in *twice htp-2* mutants. We investigated if the extended activity of DSB-2 seen in *htp-1^htp-2 HORMA C^* mutants resulted in increased recombination intermediates by monitoring RAD-51 foci, which label intermediates downstream of DSB formation and upstream of crossover designation. Numbers of RAD-51 foci were increased in *htp-1^htp-2 HORMA C^* mutants compared to *twice htp-2* and *htp-1(gk174)* mutants (Figure 3e and Figure S2b), correlating with the extended region of DSB-2 activity observed in *1^htp-2 HORMA C^* mutants and suggesting that *htp-1^htp-2 HORMA C^* mutants undergo increased DSB formation compared to *twice htp-2* and *htp-1(gk174)* mutants.

The defects in pairing, synapsis, and checkpoint control of meiotic prophase observed in *1^htp-2 HORMA C^* mutants reveal that HTP-1^HTP-2 HORMA C^ is functionally closer to HTP-2 than HTP-1. Our findings also reveal that functional differences between HTP-1 and HTP-2 are not due to differences in loading to axial elements, as increasing levels of HTP-2 loading is not sufficient to functionally replace HTP-1’s roles in pairing, synapsis, and checkpoint control of meiotic prophase.

### The safety belt region of the HORMA domain controls functional differences between HTP-1 and HTP-2

Having demonstrated that HTP-1^HTP-2 HORMA C^ is unable to provide key HTP-1 functions, we set out to further narrow down specific residues responsible for the functional differences between HTP-1 and HTP-2. The 10 amino acid substitutions present in HTP-1^HTP-2 HORMA C^ are contained in the interval between E191 and L250. A mutant strain carrying the E191A and E200K substitutions did not show obvious meiotic defects, and given the similarity between aspartic (D) and glutamic acid (E) we considered that the E215D substitution was unlikely to induce a strong effect. Thus, we focused on the 7 remaining mutations located between D226 and L250, corresponding to the safety belt of the HTP-1/2 HORMA domain, by creating mutants containing 3, 5, and 7 HTP-1-HTP-2 substitutions (Figure 4a). We found a clear additive effect when comparing these mutants, with embryonic lethality and crossover defects increasing with the number of substitutions (Figures 4b-c). Similar to *htp-1^htp-2 HORMA C^* mutants, *htp-1^mut7^* displayed 96% embryonic lethality and an average of 9.2 DAPI-stained bodies in diakinesis oocytes (Figures 4b-c). Thus, we conclude that the crossover defect initially observed in *htp-1^htp-2 HORMA C^* mutants is due to the seven amino acid substitutions present on the safety belt region. Moreover, similar to *htp-1^htp-2 HORMA C^* mutants, *htp-1^mut7^* mutants also display a strong defect in SC assembly, with pachytene nuclei displaying multiple unsynapsed regions (Figure 4d). The meiotic defects observed in *htp-1^mut7^* mutants were not due to impaired loading to axial elements, as levels of HTP-1^mut7^ protein were higher than those observed for WT HTP-1 (Figure 4e). Thus, the safety belt region of HTP-1, which is involved in the binding of HTP-1 to the closure motifs in HIM-3 and HTP-3 ^25^, is key for HTP-1 functions, but not simply by promoting chromosomal loading. These findings identify the safety belt region as key to explain the functional differences between HTP-1 and HTP-2.

**Figure 4.**
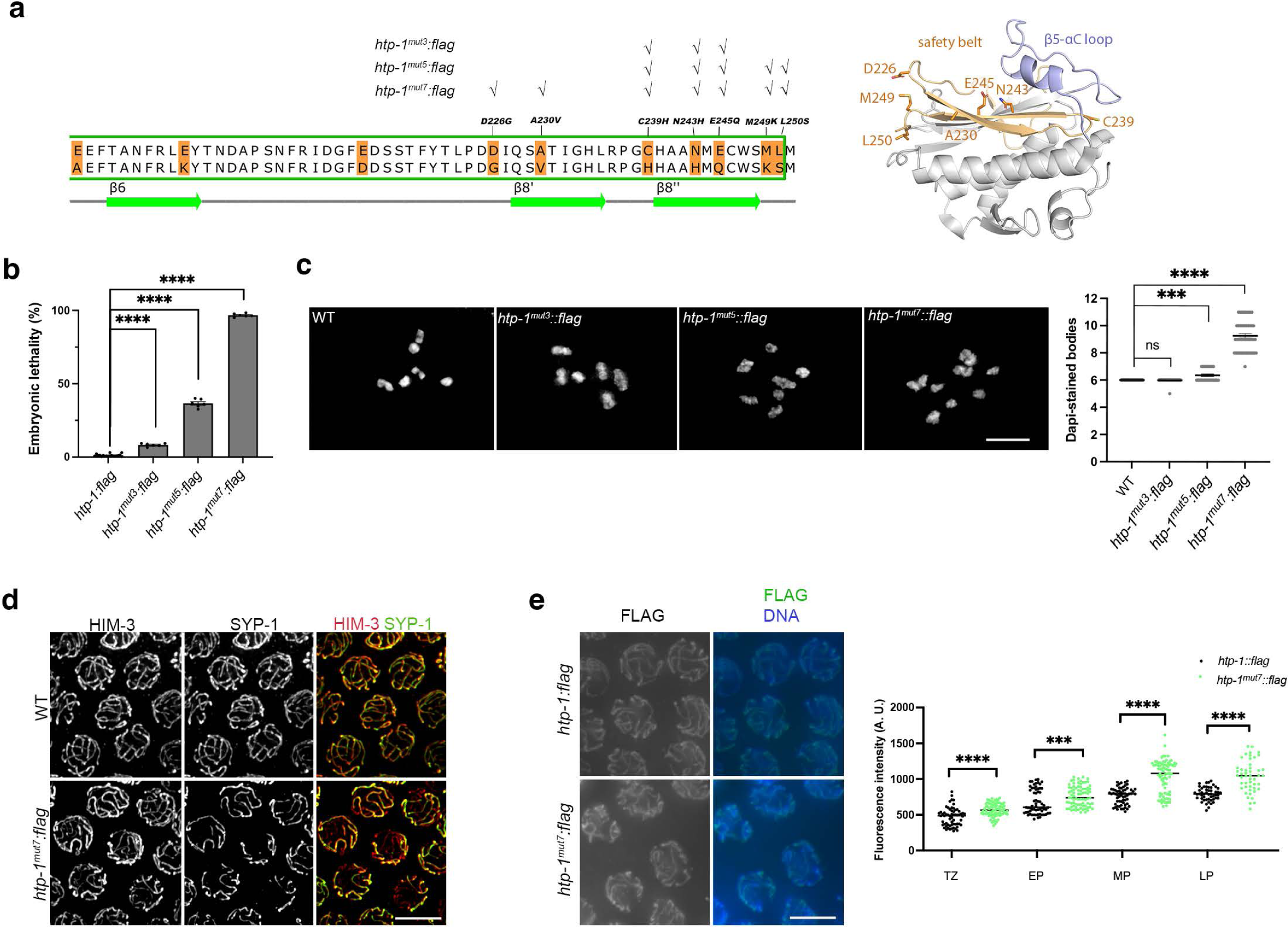
Amino acid substitutions in the safety belt region of HTP-1 induce meiotic defects. **a)** Alignment of HTP-1 and HTP-2 indicating amino acid substitutions (highlighted in orange) and secondary structure features of both proteins. Tick symbols above alignment indicate amino acid substitutions included in each HTP-1 mutant protein. Right-hand panel shows the position of the amino acid substitutions in the crystal structure of HTP-1 HORMA domain in the closed conformation (PDB 4TZQ). **b)** Quantification of embryonic lethality in strains of indicated genotypes. Number of worms and embryos analysed per genotype: *htp-1::FLAG*= 18, 5198; *htp-1^mut3^::FLAG*= 6, 1781; *htp-1^mu5^::FLAG*= 6, 1508; *htp-1^mut7^::FLAG*= 6, 871, error bars indicate mean with 95% CI, p values were calculated using a two-tailed Mann-Whitney U test. **c)** Projections of diakinesis oocytes of indicated genotypes stained with DAPI. 6 DAPI-stained bodies indicate normal crossover formation, while higher numbers indicate a crossover defect. Graph shows quantification of number of DAPI-stained bodies per genotype WT= *30*; *htp-1^mut3^::FLAG*= 50; *htp-1^mu5^::FLAG*= 36; *htp-1^mut7^::FLAG*= 48; error bars indicate mean with 95% CI, p values were calculated using a two-tailed Mann-Whitney U test. **d)** Projections of pachytene nuclei stained with anti-HIM-3 (axial element) and anti-SYP-1 (SC component) antibodies. Lines displaying only HIM-3 staining (red signal) indicate the presence of unsynapsed regions, which are only detected in *htp-1^mut7^::FLAG* mutants. **e)** Non-deconvolved projections of pachytene nuclei from indicated genotypes stained with anti-FLAG antibodies and DAPI. Images were acquired and adjusted with the same settings for both genotypes. Graphs show intensity of anti-FLAG staining in nuclei of the indicated germline regions (transition zone (TZ), early pachytene (EP), mid pachytene (MP), and late pachytene (LP) and genotypes). Between 48 and 85 nuclei from three or four different germlines were analysed per stage and genotype, error bars indicate mean with 95% CI, p values were calculated using a two-tailed Mann-Whitney U test. Note increased levels of HTP-1^mut7^::FLAG binding compared to HTP-1::FLAG. Scale bar =5 µm in all panels.

### Molecular dynamics simulations uncover different behaviour of HTP-1 and HTP-2 HORMA domains

Given that HTP-1 and HTP-2 are highly similar in protein structure and interact with the same closure motifs on HTP-3 and HIM-3 ^25^, our findings above suggest that the dynamic regulation of HTP-1/2’s HORMA domain may underscore functional differences between these highly similar paralogs. We tested this hypothesis using equilibrated molecular dynamics (MD) simulations to compare the behaviour of HTP-1, HTP-2, and HTP-1^mut7^ HORMA domain (residues 17-253) over a 500 ns interval. Simulations were based on the crystal structures of HTP-1’s (PDB 4TZQ) and HTP-2’s (PDB 4TZM) HORMA domain in the closed conformation ^25^. Simulations were run without the bound closure motifs and repeated three independent times. Over the course of the simulations, despite lacking their bound closure motifs, all three proteins retained their closed unliganded conformation (empty closed conformation from now on) (Figures 5a-b). To identify regions of greatest flexibility, we calculated the root mean square fluctuation (RMSF), which measures the average deviation of each residue from its starting position over the course of the simulation (Figure S3a). These revealed a high degree of flexibility in residues within the N-terminus, the β2-β3 hairpin, the β5-αC loop, and those within the binding pocket loop. The highest flexibility (excluding the very N and C-termini) was found in residues 140-150 corresponding to the β5-αC loop. This was also evident in the root mean square deviation (RMSD) measurements of residues 131-163, which suggested the β5-αC loop in HTP-1^mut7^ showed the greatest degree of movement from its starting position (Figure S3b). Principle component analysis of the three simulations showed that the β5-αC loop moved slightly upwards (away from the core of the of the HORMA domain) in HTP-1, while in HTP-1^mut7^ the loop moved downwards sitting on top of the safety belt formed by β8’ and β8” (Figure 5c). The loop also moved downwards in HTP-2, although in a less pronounced way than in HTP-1^mut7^ (Figures 5b-c). To quantify the movement of the β5-αC loop, we measured the distance between the centre of mass of residues in the β5-αC loop (D144 to Q148) and a group of residues on the safety belt towards which the extended loop moves downwards (D225 to I227). The mean distance between these two residue groups (measured from three independent 500 ns simulations) confirmed a significant decrease in HTP-2 (2.00 ± 0.21 nm) and HTP-1^mut7^ (1.54 ± 0.44 nm) compared to HTP-1 (2.18 ± 0.21 nm) (Figure 5d). A closer look at the simulations suggested the increased downward motion identified in the β5-αC loops of HTP-1^mut7^ and HTP-2 were likely influenced, at least in part, by the residues at positions 239 and 243 (Figure S3c-d). In the simulations of both HTP-1^mut7^ and HTP-2, the distance between the α-carbon of H239 to that of G135 fluctuated substantially compared to HTP-1 and increased from a mean distance of 0.72 ± 0.09 nm in WT HTP-1 to 0.96 ± 0.07 nm and 0.90 ± 0.06 nm in HTP-1^mut7^ and HTP-2 respectively. This difference is likely a consequence of the amino acid substitution from cysteine in HTP-1 to histidine in HTP-1^mut7^ and HTP-2 (Figure S3d). The increased flexibility at the end of β5 in HTP-2 and HTP-1^mut7^, was accompanied by an additional movement within the small α-helix at the opposite end of the β5-αC loop. In all three simulations of HTP-1^mut7^ and one simulation of HTP2, this α-helix twisted to form an alternate position (Figure S3c), which was stabilised by interactions between H243, H152 and F153.

**Figure 5.**
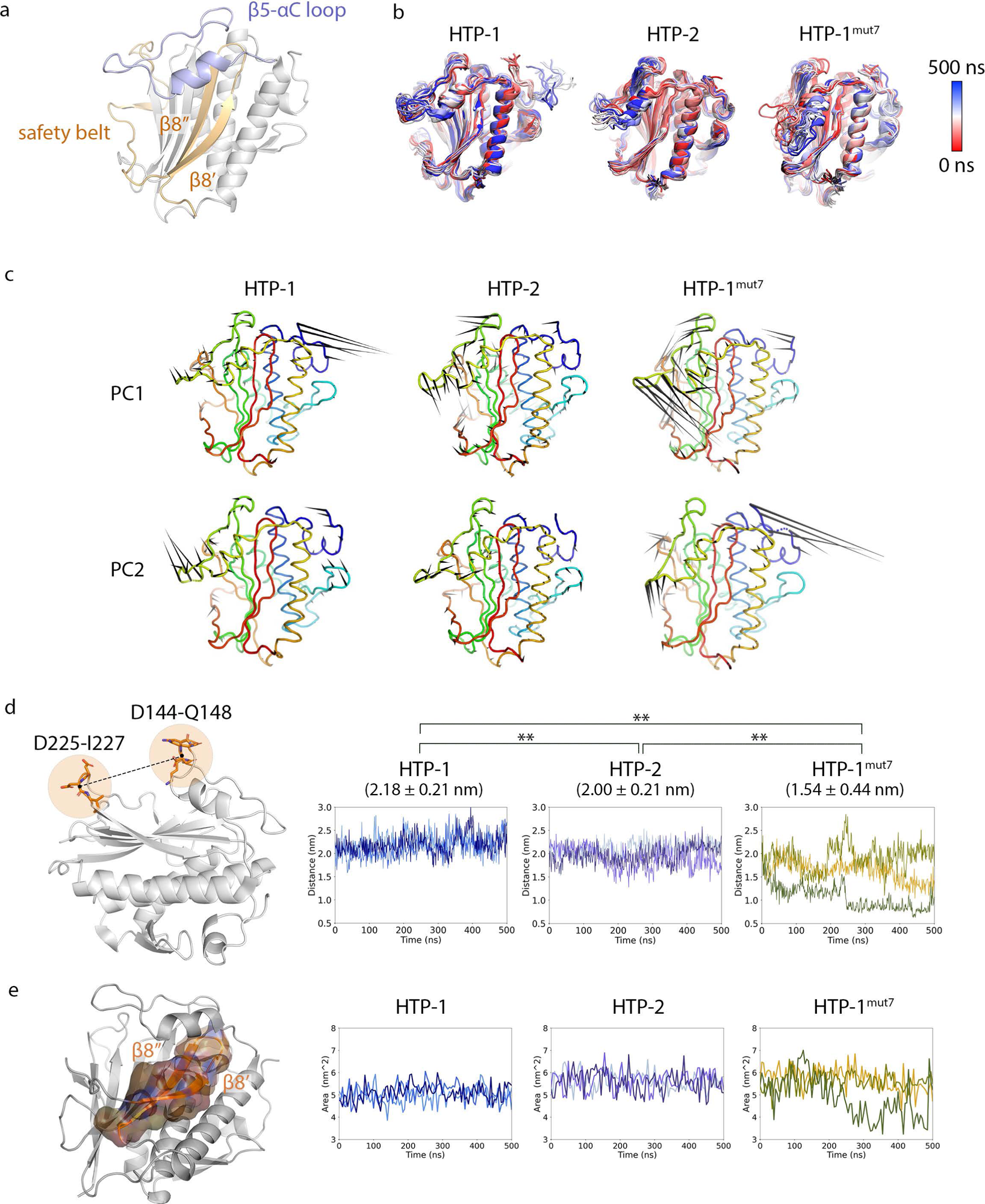
Molecular dynamics simulations of HTP-1, HTP-2, and HTP-1^mut7^ predict different dynamic behaviours. **a)** Crystal structure of HTP-1’s HORMA domain in the closed conformation (PDB 4TZQ), highlighting the β5-αC loop (blue) and safety belt (orange). **b)** protein dynamics of HTP-1 HTP-2 and HTP-1^mut7^ over the course of the 500 ns simulation. Frames every 20 ns were superimposed and aligned using protein backbone excluding residues within the β5-αC loop. **c)** Principal component analysis on triplicate 500 ns simulations. Arrows indicate the direction and magnitude of motion. The first two principal components (PC) are shown. Note that in PC1 the β5-αC loop moves downwards in HTP-2 and HTP-1^mut7^. **d)** Distance measurements of the three independent simulations between the centre of mass of D144-Q148 and D225-I227. Note the decrease in distance in HTP-1^mut7^. **e)** Solvent assessable surface area (SASA) measurements of β8’ and β8’’ in HTP-1, HTP-2 and HTP-1^mut7^ over the course of the 3 independent 500 ns simulations. Note the decrease in SASA for the β8’ and β8’’ region in HTP-1^mut7^.

The findings above suggest that the β5-αC loop is a highly dynamic region of HTP-1/2 even when the HORMA domain is in a closed conformation. Moreover, in HTP-1^mut7^ and HTP-2 simulations the downwards movement places the loop in close physical proximity to the safety belt region (Figures 5b-c), suggesting that the loop could act to stabilise the closed conformation of the HORMA domain by interfering with the conformational changes that the safety belt is expected to undergo in the unlocking of closed-conformation HTP-1/2. To determine the effects of this β5-αC loop movement on the safety belt region in HTP-1, HTP-1^mut7^, and HTP-2 we compared the solvent accessible surface area (SASA) of the safety belt (I227 to S248) in these simulations of the three proteins. This revealed a decreased SASA of HTP-1’s safety belt (average 5.06 ± 0.38 nm^2^) compared to that of HTP-2’s (average of 5.63 ± 0.44 nm^2^) (Figure 5e). Thus, in the closed conformation, the safety belt of HTP-2 is more solvent exposed, which could explain its decreased association to the axis in the presence of HTP-1. Although the SASA of HTP-1^mut7^ begins at a level comparable to that of HTP-2, the larger downward motion seen in the β5-αC loop over the course of the simulations results in a large decrease in SASA. This presumably protects the protein from release and could explain the increase abundance seen associating with the chromosome axis in vivo. In summary, the simulations of HTP-1, HTP-2, and HTP-1^mut7^ suggest that the interplay between the β5-αC loop and safety belt regions of HTP-1/2’s HORMA domain may be an important factor in determining the functions of these proteins.

### The β5-αC loop regulates HORMADs association with axial elements

Our MD simulations suggest that the β5-αC loop may play an important role in controlling the conformation of HTP-1/2’s HORMA domain. We tested the functionality of this domain in vivo by creating CRISPR mutants carrying either a whole (HTP-1^Δ140-162^) or partial (HTP-1^Δ151-162^) deletion of the β5-αC loop region (Figures 6a-b). Crossover formation was impaired in both *htp-1^Δ140-162^* and *htp-1^Δ151-162^* mutants, as evidenced by the presence of elevated numbers of DAPI-stained bodies in diakinesis oocytes (average of 11 and 9.4 respectively, compared to 6 in controls) (Figure 6c). Interestingly, differences in the number of DAPI-stained bodies between *htp-1^Δ140-162^* and *htp-1^Δ151-162^* mutants are statistically significant (Figure 6c), evidencing that HTP-1^Δ151-162^ retains some crossover-promoting activity. To further explore the functionality of HTP-1^Δ140-162^ and HTP-1^Δ151-162^, we first investigated SC assembly in *htp-1^Δ140-162^* and *htp-1^Δ151-162^* mutants. SC assembly was severely disrupted in both mutants, with almost a 100% of pachytene nuclei displaying unsynapsed regions (Figure 6d). Given this, we next tested whether HTP-1^Δ140-162^ and HTP-1^Δ151-162^ were capable of providing checkpoint control of SC assembly by delaying meiotic prophase in the presence of unsynapsed chromosomes. Staining with anti-PLK-2 antibodies revealed that *htp-1^Δ140-162^* mutants failed to accumulate nuclei with PLK-2 aggregates, while *htp-1^Δ151-162^* mutants displayed accumulation of nuclei with PLK-2 aggregates but only into the early pachytene region of the germline (Figure 6e). Thus, HTP-1-dependent checkpoint control of meiotic progression is partially functional in *htp-1^Δ151-162^* mutants, but is impaired in *htp-1^Δ140-162^* mutants. Together with the stronger crossover defect found in *htp-1^Δ140-162^* mutants, these findings suggest that *htp-1^Δ140-162^* behaves as a null allele while *htp-1^Δ151-162^* is a reduction of function allele.

**Figure 6.**
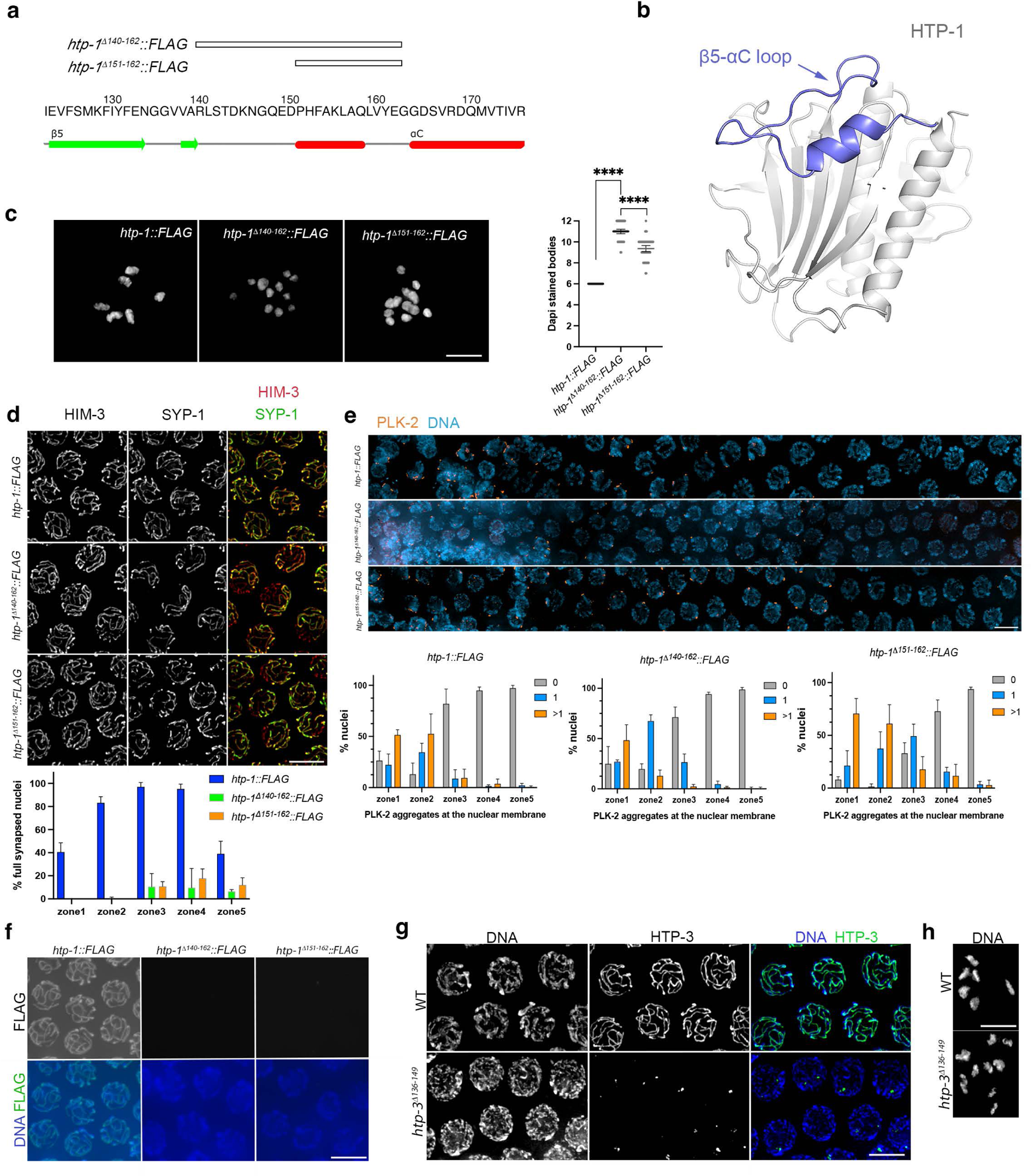
The β5-αC loop of HTP-1 and HTP-3 is required for axis loading and crossover formation. **a)** Sequence and secondary structure features of HTP-1’s β5-αC loop region indicating amino acids deleted in two HTP-1 loop mutant proteins. **b)** Closed conformation model of HTP-1 indicating the position of the β5-αC loop. **c)** Projections of diakinesis oocytes of indicated genotypes stained with DAPI. 6 DAPI-stained bodies indicate normal crossover formation, while higher numbers indicate a crossover defect. Graph shows quantification of number of DAPI-stained bodies per genotype: *htp-1::FLAG*= 40; *htp-1^Δ140-162^::FLAG*= 57; *htp-1^Δ151-162^::FLAG*= 52, error bars indicate mean with 95% CI, p values were calculated using a two-tailed Mann-Whitney U test. **d)** Projections of pachytene nuclei stained with anti-HIM-3 (axial element) and anti-SYP-1 (SC component) antibodies. Lines displaying only HIM-3 staining (red signal) indicate the presence of unsynapsed regions. Graph shows quantification of the percentage of nuclei displaying full synapsis (complete overlap of HIM-3 and SYP-1 signals) in five zones along the germline. Note the extensive presence of nuclei with unsynapsed regions in *htp-1^Δ140-162^::FLAG* and *htp-1^Δ151-162^::FLAG* mutants. Number of nuclei analysed per zone (3 germlines per genotype): 148, 150, 167, 133, 110 (*htp-1::FLAG*); 153, 148, 131, 105, 95 (*htp-1^Δ140-162^::FLAG*); 174, 187, 168, 120, 97 (*htp-1^Δ151-162^::FLAG*) mutants. **e)** Projections of nuclei in transition zone and early pachytene regions of the germline stained with anti-PLK-2 antibodies and DAPI. Nuclei with more than 1 PLK-2 aggregate indicate high CHK-2 activity, 1 PLK-2 aggregate indicates intermediate CHK-2 activity and no PLK-2 aggregate indicates no CHK-2 activity. Graphs show quantification of the % of nuclei with a given number of PLK-2 aggregates in five zones along the germline. Number of nuclei (three germlines per genotype) analysed per zone= 176, 159, 148,138, 127 (*htp-1::FLAG*); 159, 183, 183, 129, 109 (*htp-1^Δ140-162^::FLAG*); 147, 138, 130, 114, 102 (*htp-1^Δ151-162^::FLAG)*. **f)** Non-deconvolved projections of pachytene nuclei from indicated genotypes stained with anti-FLAG antibodies and DAPI. Images were acquired and adjusted with the same settings for both genotypes. Note the absence of axis associated signal in *htp-1^Δ140-162^::FLAG* and *htp-1^Δ151-162^::FLAG* mutants. **g)** Projections of pachytene nuclei of indicated genotype stained with anti-HTP-3 pS285 antibodies and DAPI. Note that HTP-3*^Δ136-149^* fails to form axial elements. **h)** Projections of diakinesis oocytes of indicated genotypes stained with DAPI. Note the presence of more than 6 DAPI-stained bodies in *htp-3^Δ136-149^*. Scale bar =5 µm in all panels.

Our MD simulations also suggest that residues within the β5-αC loop could interact with residues on the safety belt region that mediates binding of HTP-1 to closure motifs on HTP-3 and HIM-3. Thus, we evaluated the loading of HTP-1^Δ140-162^ and HTP-1^Δ151-162^ to axial elements. Strikingly, anti-FLAG staining revealed that both mutant proteins failed to load to axial elements to levels that we could detect (Figure 6f), revealing that the β5-αC loop is required for HTP-1 loading to axial elements. Despite this, HTP-1^Δ151-162^ partially supports HTP-1 checkpoint function (see above), supporting the proposal that this role involves a pool of nuclear soluble HTP-1 ^36^. As the presence of an extended β5-αC loop is a conserved feature of meiotic HORMADs, we asked if this region is also required for chromosomal loading of HORMADs beyond HTP-1. By deleting the β5-αC loop of HTP-3 we confirmed that, similar to HTP-1, deletion of the β5-αC loop in HTP-3 hindered chromosomal loading and crossover formation (Figure 6g-h). These results suggest that the β5-αC loop is required for the acquisition of a conformation that enables chromosomal loading of meiotic HORMADs.

### The β5-αC loop determines the folding conformation of meiotic HORMADs

We next considered models that could explain how the β5-αC loop may promote chromosomal loading of meiotic HORMADs. First, the β5-αC loop may be required for the conformational changes involved in the formation of an unbuckled intermediate that allows binding to closure motifs present on chromosome-bound proteins. Using ColabFold, we observed that the HORMA domain of HTP-1, HTP-2, HTP-1^Δ140-162^, HTP-1^Δ151-162^, and HTP-3^Δ136-149^ are all predicted to display an empty closed conformation (Figure 7a-b and Figure S4a-b). Similarly, a yeast Hop1 mutant protein lacking the β5-αC loop is thought to display a closed conformation in vitro ^26^. These observations are consistent with HTP-1^Δ140-162^, HTP-1^Δ151-162^, and HTP-3^Δ136-149^ being folded into a stable closed conformation that could hinder binding to closure motifs on their chromosomal interactors. As HTP-1 and HTP-3 are not thought to bind closure motifs on their own C-terminus ^25^, we refer to this conformation as “empty closed”. We tested the idea that a stably closed HTP-1 conformation would prevent chromosomal loading by substituting the last 10 amino acids of HTP-1 with HIM-3’s closure motif, as this is expected to result in a self-closed HTP-1 (Figure 7c). Similar to HTP-1 loop mutants, HTP-1^HIM-3CM^ failed to load on chromosomes, causing defects in synapsis and chiasma formation (Figure 7d-f). Interestingly, *htp-1^HIM-3CM^* mutants displayed extensive accumulation of nuclei with PLK-2 aggregates (Figure S4c), suggesting that soluble self-closed HTP-1^HIM-3CM^ is capable of delaying meiotic progression in the presence of synapsis defects and providing further support for the role of nuclear soluble HTP-1 in this process. Together, these observations support the possibility that mHORMADs loopless mutants fold into a stable closed conformation that hinders binding to closure motifs on chromosomal interactors.

**Figure 7.**
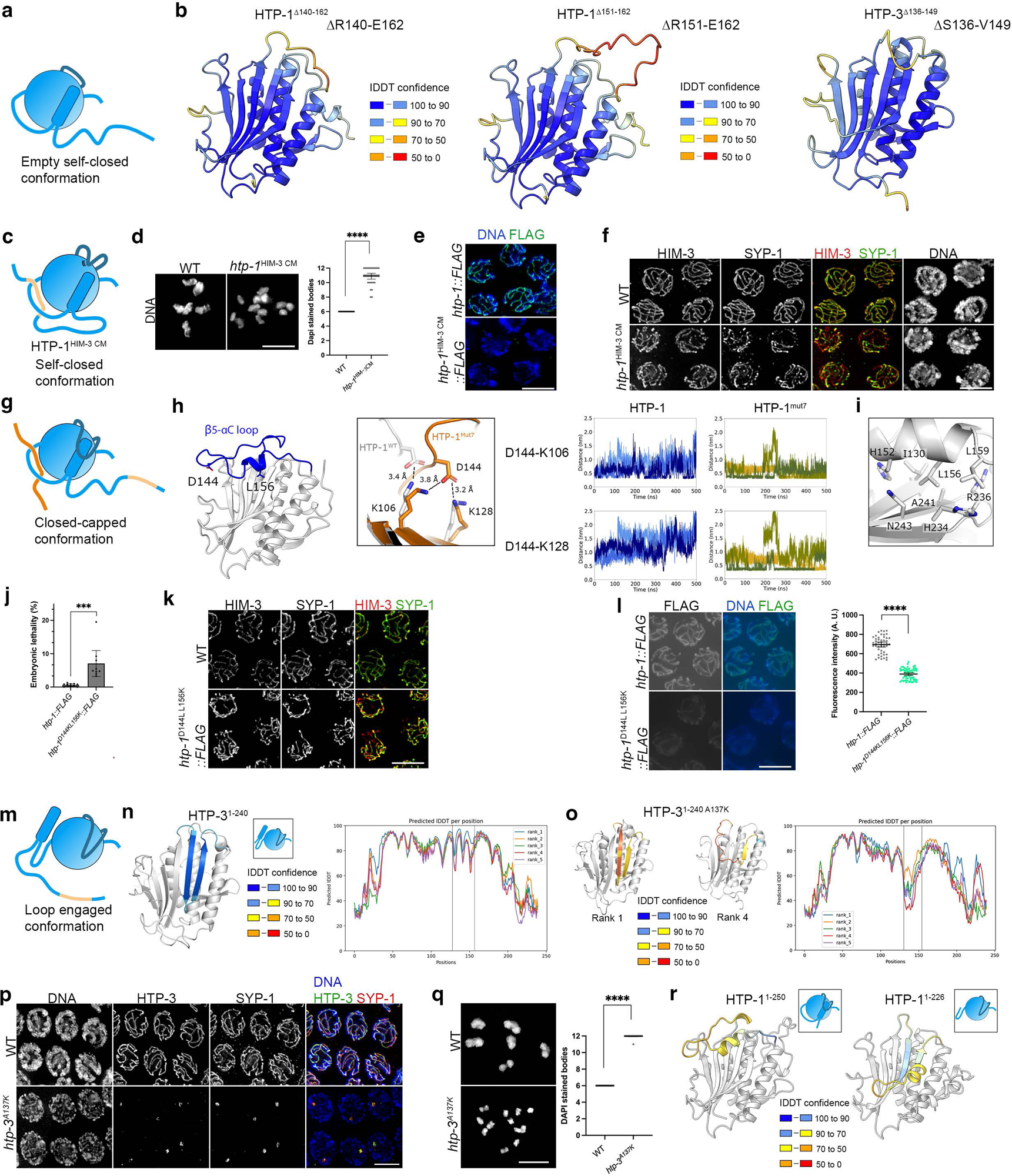
The position of the β5-αC loop and the safety belt determine the conformation of HTP-1 and HTP-3. **a)** Cartoon of mHORMAD in an empty closed conformation (circle: HORMA core, dark blue line (β5-αC loop), light blue line (safety belt region). **b)** ColabFold predictions of indicated HTP-1 and HTP-3 loop deletion mutants, which display an empty self-closed conformation. **c)** Cartoon of mHORMAD in a self-closed conformation binding to a closure motif on its C-terminal tail. **d)** Projections of diakinesis oocytes of indicated genotypes stained with DAPI. 6 DAPI-stained bodies indicate normal crossover formation, while higher numbers indicate a crossover defect. Graph shows quantification of number of DAPI-stained bodies per genotype: WT= 24; *htp-1^HIM-3 CM^*= 34, error bars indicate mean with 95% CI, p values were calculated using a two-tailed Mann-Whitney U test. **e)** Projections of pachytene nuclei of indicated genotype stained with anti-FLAG antibodies (HTP-1) and DAPI, note absence of axis staining in *htp-1^HIM-3 CM^* mutants. **f)** Projections of pachytene nuclei of indicated genotype stained with anti-HIM-3 (axis component) and anti-SYP-1 (SC component) antibodies and DAPI. Note extensive asynapsis in *htp-1^HIM-3 CM^* mutants. **g)** Cartoon of mHORMAD in a closed-capped conformation. **h)** HTP-1 structure showing the position of D144 and L156. Middle panel shows interactions between D144 and K106 or K128 in the MD simulations of WT HTP-1 and HTP-1^mut7^. Graphs show the distance between D144-K106 and D144-K128 in WT HTP-1 and in HTP-1^mut7^. **i)** HTP-1 structure displaying the position of L156 on the short helix at the C-terminus of β5-αC loop, sitting in a hydrophobic pocket. **j)** Embryonic lethality in *htp-1::FLAG* controls and *htp-1^D144K L156K^::FLAG* mutants. Number of worms and embryos analysed per genotype: *htp-1::FLAG*= 6, 1373; *htp-1^D144K L156K^::FLAG*= 9, 1840, error bars indicate mean with 95% CI, p values were calculated using a two-tailed Mann-Whitney U. **k)** Projections of pachytene nuclei stained with anti-HIM-3 (axial element) and anti-SYP-1 (SC component) antibodies. Note that unsynapsed regions are present in *htp-1^D144K L156K^::FLAG* mutants. **l)** Non-deconvolved projections of pachytene nuclei from indicated genotypes stained with anti-FLAG antibodies and DAPI. Images were acquired and adjusted with the same settings for both genotypes. Graph shows intensity of anti-FLAG staining in pachytene nuclei, note reduction of signal intensity in *htp-1^D144K L156K^::FLAG* mutants. 47 nuclei were analysed in *htp-1::FLAG* and 60 in *htp-1^D144K L156K^::FLAG*, p values were calculated using a two-tailed Mann-Whitney U test. **m)** Cartoon of mHORMAD in a loop engaged conformation. **n-o)** ColabFold predictions of indicated HTP-3 versions, indicating IDDT confidence for the β5-αC loop according to colors indicated on key. Models corresponding to rank 1 and rank 4 are shown for HTP-3^1-240 A137K^. Graphs show the IDDT confidence per position with vertical lines indicating the position of the β5-αC loop and colors indicating IDDT of five different models (ranks 1-5). Note decreased IDDT confidence for the position of the β5-αC loop in HTP-3^1-240 A137K^. **p)** Projections of pachytene nuclei of indicated genotype stained with anti-HTP-3 pS285 and anti-SYP-1 (SC component) antibodies and DAPI. Note that HTP-3^A137K^ fails to form axial elements and colicalizes with SYP-1 in nuclear aggregates. **q)** Projections of diakinesis oocytes of indicated genotypes stained with DAPI. 6 DAPI-stained bodies indicate normal crossover formation, while higher numbers indicate a crossover defect. Graph shows quantification of number of DAPI-stained bodies per genotype WT= 24; *htp-3^A137K^*= 36, error bars indicate mean with 95% CI, p values were calculated using a two-tailed Mann-Whitney U test. **r)** ColabFold predictions of indicated HTP-1 versions indicating IDDT confidence for the β5-αC loop according to colors indicated on key. Note that HTP-1^1-250^ models as an empty closed conformation while HTP-1 ^1-226^ (safety belt deletion) models as a loop engaged conformation. Scale bar =5 µm in all panels.

A second model is that the β5-αC loop may also act to stabilise the closed conformation of mHORMADs bound to closure motifs on chromosome-bound HTP-3 or HIM-3 by moving downwards towards the HORMA core, as suggested by our simulations of HTP-2 and HTP-1^mut7^ (Figure 5). We refer to this as a “closed capped” conformation (Figure 7g). To test this possibility, we examined our simulations to identify residues expected to promote or stabilise the lowered position of the β5-αC loop. In all simulations, residue D144 within the β5-αC loop is predicted to form a stable interaction with K106 on β4 (Figure 7h). In the HTP-1^mut7^ simulations, D144 also comes into close proximity to residue K128 on β5, potentially further stabilising the lowered position of the loop (Figures 7h). In addition, L156 on the short alpha helix at the C-terminal region of the β5-αC loop anchors this region to the HORMA domain by sitting in a hydrophobic pocket (Figure 7i). We reasoned that introducing a charge reversal mutant at position 144 (D114K) and introducing a charge at position 156 (L156K), would disrupt these interactions and interfere with the downwards movement of the β5-αC loop towards the safety belt. Thus, we used CRISPR to generate an *htp-1^D144K L156K^* allele to investigate the effect of these mutations in vivo. *htp-1^D144K L156K^* mutants displayed increased levels of embryonic lethality, synapsis defects, and the accumulation of nuclei with PLK-2 aggregates in the pachytene region (Figures 7j-k and Figure S4d). Next, we monitored the loading of HTP-1^D144K L156K^::FLAG to meiotic chromosomes using anti-FLAG antibodies. The amount of HTP-1^D144K L156K^::FLAG associated with axial elements of pachytene nuclei showed a significant reduction compared to WT HTP-1::FLAG controls (Figure 7l). These findings confirm that the β5-αC loop region modulates the loading and function of HTP-1 and suggest that it may do so, at least in part, by interacting with residues in the HORMA core as suggested by the “capping” model.

We also used ColabFold to predict the structure of HTP-3’s HORMA domain to determine if, similar to HTP-1, the β5-αC loop of HTP-3 could also cap the safety belt region. Strikingly, the β5-αC loop of HTP-3 is predicted to fold down over the backbone of the HORMA domain forming hydrogen bonds with residues in β5 and αC (Figure 7n and Fig S4d). In this novel HORMAD configuration, which we refer to as “loop engaged” (figure 7m), the position of the two-stranded beta sheets formed by the β5-αC loop is highly reminiscent of the position of the safety belt in a canonical closed conformation. Looking at this predicted structure, we assessed that introducing a charge at position A137 could destabilise the interaction between the loop and the HORMA core. This possibility was supported by the predicted structure of the HTP-3’s HORMA carrying the A137K substitution, which caused a strong decrease in the confidence of the predicted position of the loop (Figure 7o). Thus, we generated *htp-3^A137K^* homozygous mutants to investigate the impact of the A137K mutation in vivo. HTP-3^A137K^ fails to load to chromosomes causing a complete failure in synapsis and chiasma formation (Figure 7p-q), as expected when HTP-3 is not loaded to axial elements ^12^. HTP-3^A137K^ is detected in nuclear aggregates colocalizing with SC component SYP-1 (Figure 7p), suggesting that HTP-3^A137K^ is present in the nucleus but unable to form axial elements. These results reveal that the β5-αC loop is also essential for HTP-3 loading to axial elements and suggest that the acquisition of a loop engaged conformation is an important aspect of this process.

Our findings with HTP-3 led us to ask whether other HORMADs may also adopt a loop engaged conformation. The predicted structures of HTP-1, HTP-2, HIM-3, yeast Hop1, mouse HORMAD1, and Arabidopsis ASY1 HORMA domains all adopted an empty-closed conformation, with the C-terminal region of the safety belt folded into two-stranded beta sheets that interact with β5 (Figure 7r and Figure S4a,e-h). However, in the loop engaged conformation of HTP-3 the β5-αC loop takes a very similar position to that of the two-stranded beta sheets formed by the safety belt in a canonical mHORMAD closed conformation, suggesting that the safety belt and the β5-αC loop may compete to bind to the same region of the HORMA core. Thus, we used ColabFold to predict if deleting the safety belt region of HTP-1, HIM-3, Hop1, HORMAD1, and ASY1 would result in an interaction between the β5-αC loop and the HORMA core. Indeed, deletion of the safety belt region in these mHORMADs resulted in the movement downwards of the β5-αC loop to form the “loop engaged” conformation (Figure 7r and Figure S4e-h). According to these predictions, acquisition of the loop engaged conformation involves refolding and repositioning of a flexible protein region, evidencing clear parallels with the changes that the safety region undergoes in the transition from open to closed Mad2. These findings suggest that the ability of the flexible β5-αC loop to acquire secondary structure and interact with the HORMA core is a conserved feature of mHORMADs.

## Discussion

Our findings suggest that mHORMADs have expanded the conformation landscape of their HORMA domain beyond the canonical open and closed conformations found in Mad2, with the extended β5-αC loop playing a key role in this process. Our efforts to determine the mechanistic basis for the functional differences between mHORMAD paralogs HTP-1 and HTP-2 show that these are due to 7 amino acid substitutions within the safety belt region of the HORMA domain, and not to differences in time of expression or amount of protein associated with axial elements. By combining functional in vivo studies with molecular dynamics modelling and ColabFold predictions we reveal that the interplay between two structurally flexible regions, the β5-αC loop and the safety belt, which can bind to the same region of the HORMA core are key in determining the conformation and function of these proteins. We clarify how paralogs that display nearly identical crystal structures have acquired different functions in vivo and provide evidence that the role of the β5-αC loop in controlling protein conformation and function is a conserved feature of mHORMADs. Moreover, a recent preprint proposes that the short β5-αC loop of Mad2 is also a dynamic region that modulates the open to closed Mad2 conversion ^49^, suggesting that the role of the β5-αC loop in regulating conformation dynamics is a conserved, and previously unknown, aspect of HORMAD proteins.

The model depicted in Figure 8 summarises how three different positions of the flexible β5-αC loop can lead to the formation of canonical and non-canonical mHORMAD conformations. These include: 1) A loop engaged conformation in which the β5-αC loop moves downwards to interact with β5 on the HORMA core, precluding the safety belt from interacting with this region and therefore interfering with the formation of a canonical closed conformation. Disengaging the β5-αC loop from the HORMA core could lead to the acquisition of the previously proposed funbuckled conformation ^26^, which can bind a closure motif to form a closed (closure motif bound) conformation. Thus, the loop engaged conformation may act to stabilise a conformation competent to bind closure motifs. Strong support for the functional relevance of the loop engaged configuration comes from the observation that a single amino acid substitution (A137K) on the β5-αC loop of HTP-3, which is predicted to prevent the interaction of the loop with the HORMA core, blocks axis loading of HTP-3. As axis loading is also blocked in *htp-1* and *htp-3* mutants lacking the β5-αC loop, we propose that when the β5-αC loop is prevented from binding β5 on the HORMA core mHORMADs acquire an unbound closed conformation that prevents their loading to chromosomes. 2) A closed capped conformation in which the β5-αC loop moves downwards towards the safety belt and HORMA core. Our finding that mutations predicted to weaken the interaction between the β5-αC loop and the HORMA core reduce the amount of HTP-1 bound to chromosomes suggest that this interaction may be important to stabilise the closed conformation bound to a closure motif. 3) A closed uncapped conformation in which the loop is in an upwards position without interacting with the safety belt. This conformation may represent an intermediate between the closed-capped and unbuckled conformations during the process of binding or releasing closure motifs on mHORMAD interactors. As depicted in Figure 8, closed conformations could be formed in cis, by binding to closure motifs present on the C-terminus of the protein, in trans, by binding closure motifs present in other proteins, or even without binding to a closure motif (empty closed). The existence of the empty closed conformation is also supported by the fact that Mad2 ^50^, Rev7 ^21^, and Hop1 ^26^ can all be purified as empty closed monomers.

**Figure 8.**
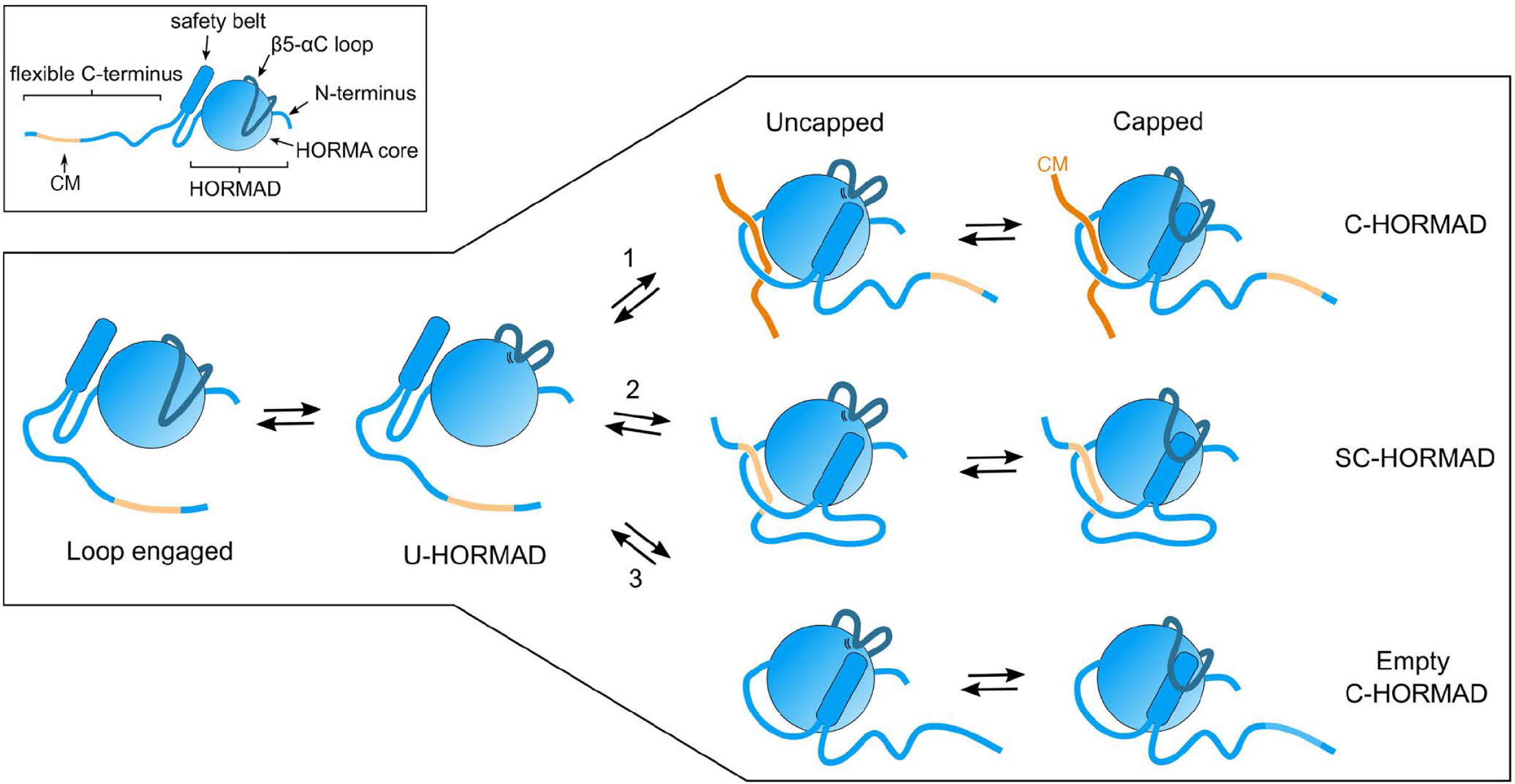
Model: The position of the safety belt and the β5-αC loop determine mHORMAD conformations. See discussion for full description. Abbreviations: CM (Closure motif), U-HORMAD (unbuckled HORMAD), C-HORMAD (closed HORMAD), SC-HORMAD (self-closed HORMAD).

Our findings help to clarify why, in contrast to Mad2 ^50^ and Rev7 ^21^, a stable open conformation has not been identified for mHORMADs ^18,26^. We propose that this is due to the presence of a second structurally flexible region, the extended β5-αC loop, which is much shorter in Mad2 and Rev7, that can associate with the same HORMA core region as the safety belt. Structural prediction of mHORMAD mutants from different organisms lacking either the safety belt or the β5-αC loop suggests that these two regions compete to bind the same HORMA core region. For example, *C. elegans* HTP-3 HORMA domain is predicted to adopt a loop engaged conformation with the β5-αC loop interacting with β5 while the safety belt is positioned to the left of β4 on the HORMA core, but deleting the β5-αC loop results in movement and refolding of the safety belt to interact with β5, thereby forming an empty closed conformation. Conversely, the predicted structures of *C. elegans* HTP-1, HTP-2 and HIM-3, yeast Hop1, *Arabidopsis* Asy1, and human HORMAD1 all adopted an empty closed conformations in which the safety belt interacts with β5 and the β5-αC loop is largely unstructured in an upwards position. In all these proteins, deletion of the safety belt leads to the prediction of the loop engaged conformation maintained by interactions between the refolded β5-αC loop and β5. Thus, the β5-αC loop and the safety belt affect the position of each other and mHORMADs appear to favour conformations in which either of these domains interacts with the HORMA core. We propose that the presence of two structurally mobile elements that can be refolded to interact with the same region of the immobile HORMA core is a conserved feature of mHORMADs that has expanded the conformational and functional complexity of these proteins.

The finding that the lexicon of mHORMADs is not limited to the previously described closed ^25,26^ (bound to a closure motif) and unbuckled ^26^ conformations provides an important milestone to elucidate the mechanisms that endow mHORMADS with the ability to control multiple meiotic events. For example, axis-bound mHORMADs are thought to orchestrate quality control of meiotic progression by sensing specific chromosomal states, such as recombination-dependent SC stability ^44^, to generate a signal that feeds back on the activity of master meiotic kinases ^37,44,51,52^. However, as axis-bound mHORMADs are expected to display a closed conformation bound to a closure motif ^25,26^, it remained unclear if this population of mHORMADs could undergo further morphological changes that may be required to start a signal in response to changes in their local chromosomal environment. Similarly, recent findings suggest that chromosome-bound Hop1 exists in two states, one that promotes DSB formation and is refractory to Pch2 removal and a second that is susceptible to Pch2 and does not promote DSB formation ^23^. The nature of these two postulated Hop1 states remains unknown. Our finding that the β5-αC loop remains as a highly flexible region even when HORMADs are in a closed conformation suggest that axis-bound mHORMADs retain conformational plasticity. For example, changes in the position of the β5-αC loop of axis-bound HORMADs could change the ability of the loop to act as an interacting surface and expose or hide other interactor-recruiting surfaces. As we have shown that the β5-αC loop of HTP-1 can move downwards to interact with the safety belt, this region is an obvious surface which availability could be regulated by the β5-αC loop. A precedent for this type of interaction is found in closed-conformation human Rev7, where the safety belt provides an interaction surface for Rev1 ^53^. In addition, both the position of the β5-αC loop and the recruitment of mHORMAD interactors to the loop or to loop-regulated surfaces are likely to be controlled by posttranslational modifications, as HORMADs are known phosphorylation and sumoylation substrates ^27,29,54,55^.

Beyond affecting the function of axis-associated mHORMADs, we provide evidence that the loop also controls the activities of nuclear soluble mHORMADS, which are proposed to participate in signal transmission of meiotic checkpoints ^36^. We have created three different HTP-1 mutants that are not detected at axial elements, thereby causing severe synapsis defects that under normal conditions are expected to trigger a HTP-1-dependent checkpoint response that sustains high levels of CHK-2 activity. First, full deletion of the β5-αC loop (HTP-1^Δ140–162^) induces a *htp-1* null-like phenotype where high levels of CHK-2 activity are not sustained despite extensive asynapsis; second, a partial loop deletion (HTP-1^Δ151-162^) results in the maintenance of intermediate levels of CHK-2 activity; and third, a self-closed version of HTP-1 (HTP-1^HIM-3 CM^) that carries an intact β5-αC loop sustains high levels of CHK-2 activity. Thus, the ability of nuclear soluble HTP-1 to sustain CHK-2 activity in the presence of synapsis defects is dictated by the integrity of the β5-αC loop. Moreover, our functional and MD comparisons between HTP-1 and HTP-2, which identified the β5-αC loop as the most dynamic region when both proteins are in a closed conformation, provide further support for the involvement of the loop in the checkpoint response to synapsis defects. In yeast, levels of Hop1 are known to modulate checkpoint responses to synapsis defects ^56,57^, suggesting that differences in the amount of HTP-1 and HTP-2 may determine their checkpoint activity. However, we show that replacing the endogenous *htp-1* locus with *htp-2* causes high levels of axis-bound HTP-2, but fails to induce an HTP-1-like checkpoint response in the absence of synapsis. Thus, we support that differences in checkpoint competence between HTP-1 and HTP-2 are due to differences in the conformational dynamics of their HORMA domains, and that the β5-αC loop plays a crucial role in this process. Future studies should address whether defects in synapsis or recombination that trigger HTP-1-dependent checkpoint responses, do so by mechanisms involving specific HORMA domain conformations.

In summary, the presence of two flexible protein regions, the β5-αC loop and the safety belt, has expanded the folding complexity of mHORMADs compared to Mad2 or Rev7. We propose that this increased folding complexity, together with the presence of N- and C-terminal domains flanking the HORMA domain endows mHORMADs with the ability to orchestrate many different aspects of meiotic chromosome metabolism, including homologue pairing, recombination, checkpoint control, and release of sister chromatid cohesion. Elucidating how different mHORMAD conformations support specific meiotic events to ensure the formation of haploid gametes remains an important and open question.

## Methods

### C. elegans genetics

All *C. elegans* strains were grown on NG agar plates seeded with OP-50 *E. coli* at 20°C. The N2 Bristol strain was used as the WT reference strain. Transgenic strains containing single-copy transgene insertions of CRISPR-mediated edits were created as described below. Table S1 contains the genotype of all strains used in this study.

Strains carrying single copy transgene insertions were created by microinjecting a strain carrying the MosCI transposon at the *ttTi5605* (chromosome II) site ^58^. Transgenes expressing chimeric HTP-1/HTP-2 proteins were generated by swapping coding regions in *htp-1* with the corresponding *htp-2* sequence. The amino acid sequence included in each domain is indicated by the first and last amino acid of that domain: N-terminus domain (1-30aa), C-terminus domain (251-352 aa), HORMA domain (31-250aa), HORMA A (31-90aa), HORMA B (91-165aa) and HORMA C (166-250aa). Table S2 contains a list of transgenes used in this study.

CRISPR-Cas9 mediated genome editing was performed using Cas9 ribonucleoprotein complexes and the co-conversion method using co-editing of *dpy-10* locus as a marker to identify edits ^59^.

### Immunostaining and Fluorescence In Situ Hybridization (FISH)

FISH and immunostaining protocols used in this study were previously described ^36^. Briefly, immunostaining was performed in germlines dissected from 19-24 hours post L4 worms in PBS buffer or EGG buffer (118 mM NaCl, 48 mM KCl_2_, 2 mM CaCl_2,_ 2 mM MgCl_2_, 5 mM HEPES) containing 0.1% Tween and immediately fixed with 1% paraformaldehyde for 5 minutes. Slides were then snap frozen in liquid nitrogen and following removal of the coverslip, slides were immersed in methanol at -20 °C for at least 1 minute. Slides were then washed three times in PBST (1xPBS, 0.1% Tween) for 5 minutes and blocked in 0.7% BSA in PBST for 30 minutes. Primary antibodies were incubated overnight at room temperature or in the cold room for RAD-51 antibodies. After three washes of 10 minutes in PBST, slides were incubated with secondaries antibodies for 2 hours in PBST at room temperature in the dark. Following three washes with PBST, slides were counterstaining with DAPI (2 μg/ml), washed with PBST for 1 hour and mounted using Vectashield (Vector). Primary antibodies used in this study: mouse anti-FLAG (M2 monoclonal F1804, Sigma), rabbit anti-PLK-2 ^60^, rabbit anti-RAD-51 ^61^, chicken anti-SYP-1 ^36^, rabbit anti-HIM-3 ^9^, rabbit DSB-2 ^42^. Secondary antibodies used in this study: goat anti-rabbit Alexa488-conjugated (Life Technologies), goat anti-chicken Alexa555-conjugated (Life Technologies), and goat anti-mouse Alexa488-conjugated (Life Technologies).

For FISH, germlines were dissected and processed as above, but using 3.7% instead of 1% paraformaldehyde. Following incubation in methanol at -20 °C, slides were washed three times (5 minutes each) in 2X SSCT (2X SSC 0.1% Tween). Slides were then dehydrated in a series of 70%, 90% and 100% ethanol for three minutes each and left to dry. The hybridization mix contained 250 ng of labelled probe (5S rDNA locus on chromosome V) in 2 x SSCT, 50 % formamide, 10% w/v dextran sulfate. FISH probes were made by labelling 1 μg of DNA with BIO-nick translation kit or DIG-nick translation kit (Roche). Following addition of hybridization mix slides were incubated in a thermocycler with the following program: 3 minutes at 93 °C, 2 minutes at 72 °C and overnight at 37 °C. Two post-hybridization washes were carried out in 2X SSCT 50% formamide at 37 °C, followed by three 5 minutes washes in 2X SSCT. Slides were then blocked in 1% BSA 2X SSCT for 1 hour and FITC-conjugated streptoavidin (Molecular Probes) and Rhodamine-conjugated anti-digoxigenin antibodies (Roche) were added in a 1:100 dilution for 1 hour at room temperature in the dark. Slides were finally washed for 30 minutes and DNA was counterstained with 1ng/ml DAPI.

Immunostaining and FISH images were acquired as three-dimensional stacks on a Delta Vision system equipped with an Olympus 1x70 microscope using a x100 lens. Images were deconvolved using SoftWoRx 3.0 (Applied Precision) and mounted using Adobe Photoshop.

### Quantification of DAPI-stained chromatin bodies in diakinesis oocytes

Germlines were dissected and stained with DAPI as described in the immunostaining protocol above. The number of DAPI-stained bodies was scored in the two most proximal oocytes (known as -1 and -2) as the level of chromosome condensation in these late diakinesis oocytes facilitates scoring of chromatin bodies.

### Quantification of RAD-51 foci in dissected germlines

Quantification of RAD-51 foci was performed in at least three germlines per genotype. Germlines were divided in six-equal size zones between the start of transition zone and the end of late pachytene and the number of RAD-51 foci were scored per nucleus in each zone. Number of RAD-51 were plotted according the following categories: 0, 1, 2 to 3, 4 to 6, 7 to 12, more than 12, and stretches (agglomerates of RAD-51 signal that were larger than individual rounded foci).

### Quantification of chromosome V FISH signals

FISH signals were quantified by dividing germline into six zones of equal length from the start of transition zone to the end of late pachytene and scoring the number of signals per nucleus. FISH signals were considered paired if a single focus was present in the nucleus and unpaired when two foci were observed in the same nucleus at a distance larger than 0.6 μm (measured in softWoRx Explorer).

### Quantification of SC assembly in dissected germlines

Germlines stained with anti-HIM-3 and anti-SYP-1 antibodies were divided into five zones of equal length from the start of transition zone until the end of late pachytene. HIM-3 and SYP-1 signals were evaluated in individual nuclei of each zone and were scored as synapsed when HIM-3 and SYP-1 signals overlapped along the full length of all chromosomes, while nuclei displaying HIM-3 regions devoid of SYP-1 signal were scored as unsynapsed.

### Quantification of PLK-2 and DSB-2 signals in dissected germlines

Germlines stained with anti-PLK-2 antibodies were divided into five equal-size zones spanning from the start of transition to the end of late pachytene. The number of PLK-2 aggregates per nucleus in each zone was scored as according to the following categories: 0, 1, or more than 1. Quantification of the DSB-2 positive zone of the germline in different genotypes was performed as previously described ^42^. Briefly, projections of germlines stained with anti-DSB-2 antibodies and DAPI were divided into vertical rows of nuclei between the onset of transition zone and the end of late pachytene. The extent of the DSB-2 positive zone was determined as the percentage of continuous rows of nuclei in which all or most nuclei displayed DSB-2 staining out of the total rows of nuclei between transition zone and the end of pachytene.

### Quantification of fluorescence signal intensities to measure HTP-1 and HTP-2

Whole-nucleus mean fluorescence intensity was used to compare the fluorescence intensity of anti-FLAG signal in dissected germlines from worms expressing different versions of *htp-1*, and *htp-2* all containing a FLAG just before their STOP codon. Images were acquired on a Delta vision microscope as 3D stacks using the same exposure settings between genotypes to be compared. Quantitative analysis of fluorescence intensity was performed as previously described ^62^. Briefly, non-deconvolved images were analysed in Image J using a customized macro. Nuclei were manually circled using the “oval” tool, one nucleus at a time, and the fluorescence intensity of each z-stack slice was measured. The mean fluorescence, displayed as arbitrary units, was calculated after normalising for the number of z-stack slices and the area drawn

### Quantification of embryonic viability

A single L4 stage hermaphrodite was placed in a plate containing a small drop of OP-50 bacteria and worms were transferred onto new plates every 12 hours. 24 hours later the number of unhatched embryos was scored from each plate. Three days after the parental worm was removed, the total number of hermaphrodite and male progeny reaching L4/adult stages were scored per plate. Embryonic viability of each biological replicate (at least 5 individual worms were used per genotype) was calculated as the number of hatched eggs over the total number of laid eggs.

### Protein structure prediction methods

Protein structural predictions were performed using ColabFold v1.5.5 AF2 with MMSeqs2. Supposition of different models were performed using PyMOL vX while figures in this manuscript were created using either PyMOL vX or ChimeraX v1.6.1.

### Computational modelling and simulation of HTP-1 and HTP-2

The crystal structures of HTP-1 (PDB 4TZQ) and HTP-2 (PDB 4TZM) ^25^ were completed by adding the missing residues (residues 6-15, 253) using Modeller 9.02 ^63^. Residue 84 in HTP-1, which is modified to leucine in the crystal structure, was replaced with proline found in the WT protein using PyMOL. HTP-1^mut7^ was generated by introducing mutations in WT HTP-1 (D226G, A230V, C239H, N243H, E245Q, M249K, L250S) using PyMOL.

The proteins were solvated by the TIP3P water model and neutralising counterions ^64^. Simulations were performed using GROMACS 2018.3 ^65^ and the CHARMM36m force field ^66^. The protein systems were firstly energy minimised using the steepest descent algorithm to relax steric clashes generated during set up. Following this, the systems were equilibrated for a total of 600 ps; the positional restraints (with force constant 1000 kJ mol^−1^ nm^−1^) initially applied to the protein backbone atoms were reduced in a stepwise manner ^67^. The production simulations of the unrestrained protein were run for 500 ns. For each system, replicate simulations were initiated using coordinates extracted at random time points from the last 100 ps of the equilibration run. The initial coordinates and velocities differ for each replicate simulation.

Simulations were performed using the NPT ensemble, with the pressure maintained semi-isotropically using the Parrinello−Rahman barostat at 1 bar and a time constant of 5 ps ^68^, and the temperature sustained at 310 K using the velocity-rescale thermostat and a coupling constant of 0.1 ps ^69^. The lengths of all bonds were constrained using the LINCS algorithm enabling a timestep of 2 fs ^70^. The Particle Mesh Ewald method was used to treat long-range electrostatic interactions with a short-range cut-off of 1.4 nm ^71^. The van der Waals interactions were curtailed at 1.4 nm, with long-range dispersion corrections applied to the pressure and energy. The periodic boundary conditions were applied to all systems in three dimensions, as done previously ^67^.

Analyses were performed using GROMACS utilities and locally written code. The molecular graphics images were generated using Visual Molecular Dynamics (VMD) package ^72^ and PyMOL. The porcupine plots for the Principal Components Analysis were generated in PyMOL. The unpaired t-test was used to assess the significance of the differences in the distances plotted in Figure 5d; *p* < 0.05 was regarded as statistically significant.

