## Supplemental Figures for "Two structurally mobile regions control the conformation and function of metamorphic meiotic HORMAD proteins"

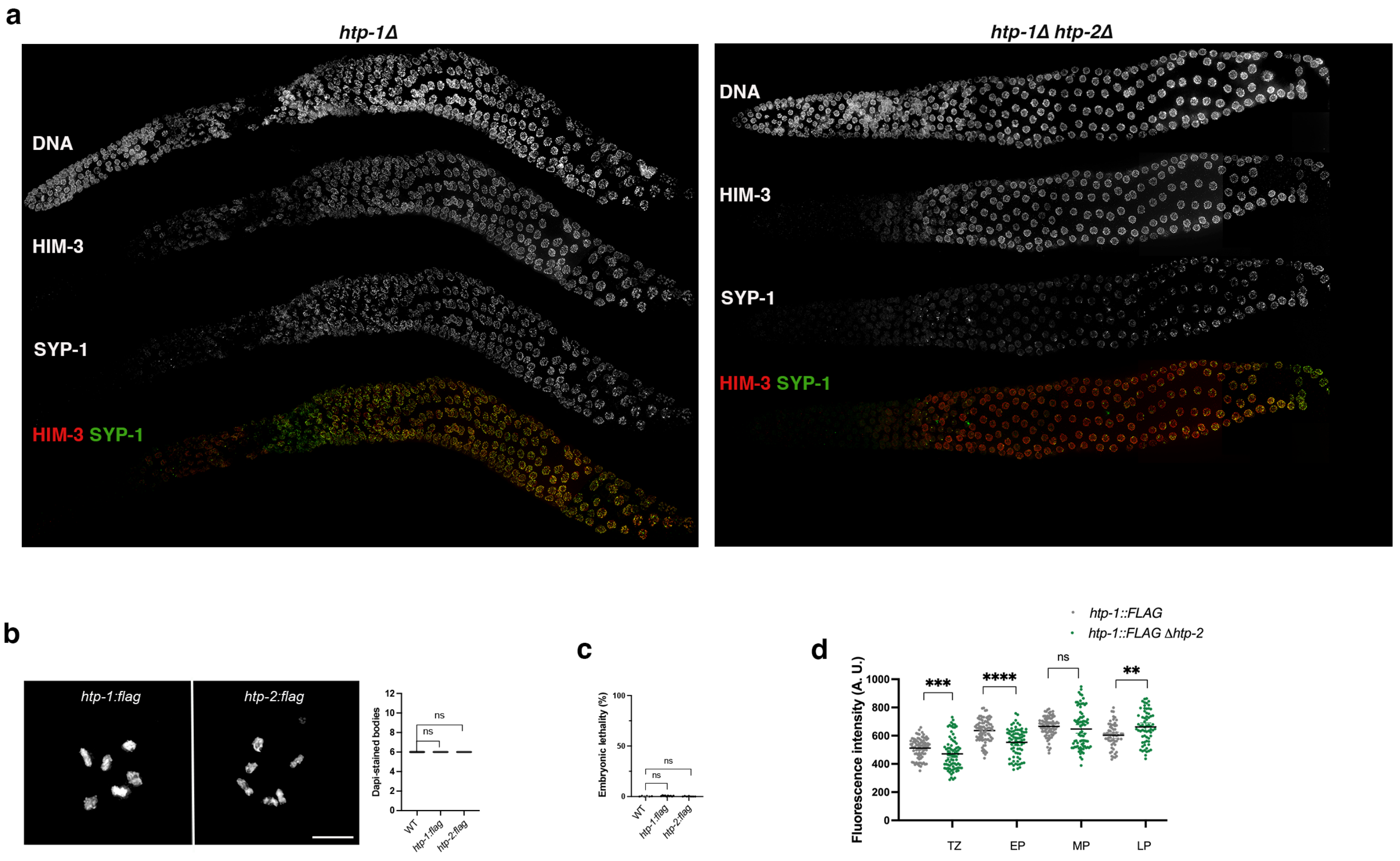


**Figure S1. a)** Projections of whole-mounted germlines of indicated genotypes stained with anti-HIM3 and anti-SYP-1 antibodies and DAPI. Note that SYP-1 staining is delayed and reduced in *htp-1Δ htp-2Δ* double mutants compared to *htp-1Δ* single mutants. **b)** Projections of diakinesis oocytes from *htp-1::FLAG* and *htp-2::FLAG* homozygous worms stained with DAPI. Note the presence of 6 DAPI-stained bodies, indicating normal crossover formation. Graph shows quantification of number of DAPI-stained bodies per genotype, between 20 and 26 oocytes were analysed per genotype, error bars indicate mean with 95% CI, p values were calculated using a two-tailed Mann-Whitney U test. Scale bar =5 µm. **c)** Quantification of embryonic lethality in strains of indicated genotypes. Number of worms and embryos analysed per genotype: WT= 6, 1707; *htp-1::FLAG*= 6, 1373; *htp-2::FLAG*= 7, 2177; error bars indicate mean with 95% CI, p values were calculated using a two-tailed Mann-Whitney U test. **d)** Graphs show intensity of anti-FLAG staining in nuclei of the indicated germline regions (transition zone (TZ), early pachytene (EP); mid pachytene (MP), and late pachytene (LP) and genotypes), between 61 and 83 nuclei from four different germlines were analysed per stage and genotype, error bars indicate mean with 95% CI, p values were calculated using a two-tailed Mann-Whitney U test.

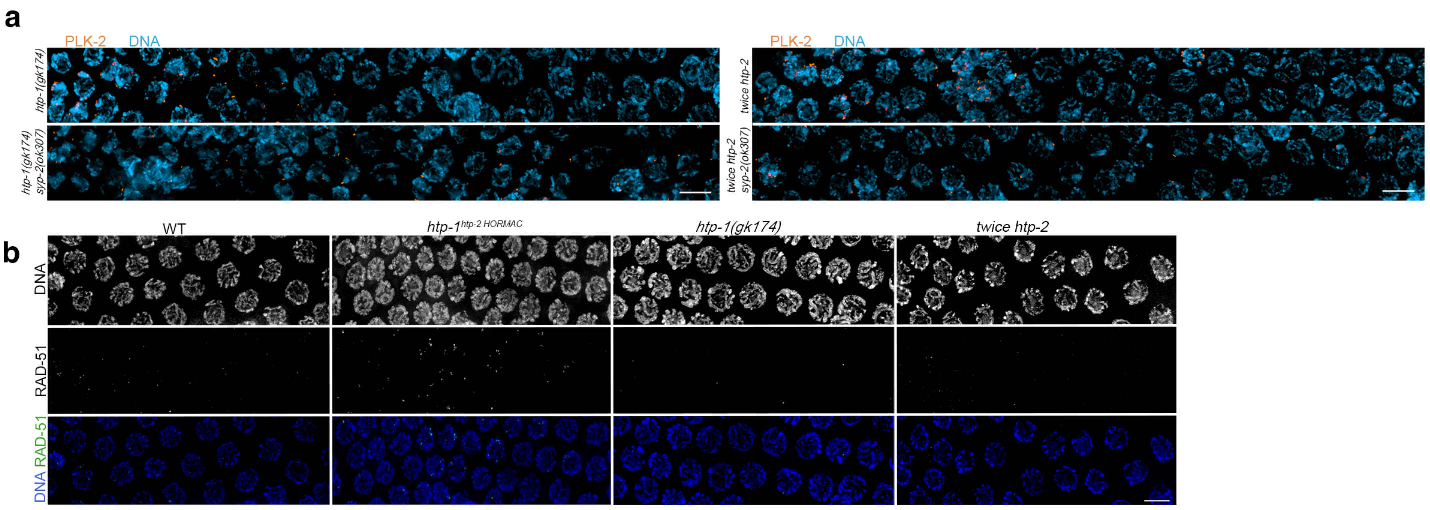


**Figure S2. a)** Projections of nuclei in transition zone and early pachytene regions of the germline of indicated genotypes stained with anti-PLK-2 antibodies and DAPI (examples of graphs shown in Fig 3c). Nuclei with more than 1 PLK-2 aggregate indicate high CHK-2 activity, 1 PLK-2 aggregate indicates intermediate CHK-2 activity and no PLK-2 aggregate indicates no CHK-2 activity. Note that in all genotypes nuclei with multiple PLK-2 aggregates are lacking in nuclei on the right-hand side of the panels, corresponding to early pachytene. **b)** Projections of pachytene nuclei (corresponding to zones 4 and 5 of the graphs shown in figure 3e) from germlines of indicated genotypes stained with anti-RAD-51 antibodies and DAPI. Scale bar =5 µm in all panels.

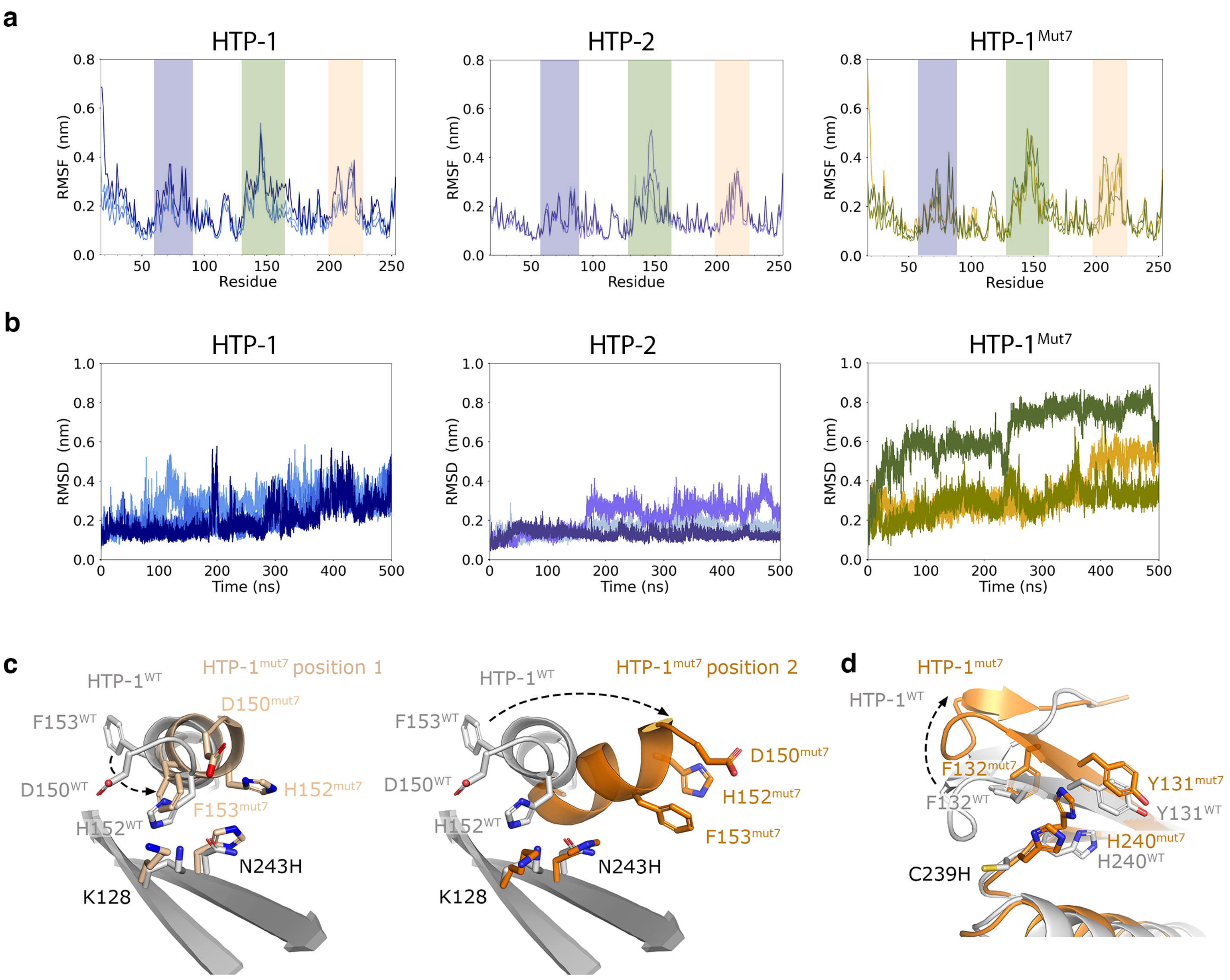


**Figure S3. a)** Root mean square fluctuation (RMSF) analysis of 3 independent simulations of HTP-1, HTP-2 and HTP-1$^{mut7}$. Residues highlighted correspond to the β2- β3 hairpin in blue, β5-αC loop in green and safety belt loop in orange. **b)** Root mean square deviation (RMSD) of the β5-αC loop (residues 131-163) in 3 independent simulations of HTP-1, HTP-2 and HTP-1mut7. **c)** Structural alignments of HTP-1 WT (grey) and HTP-1$^{mut7}$ (orange) highlighting the movements observed in the β5-αC loop. During the HTP-1$^{mut7}$ simulations 2 different positions were identified (light orange and dark orange). The alternate positions facilitated interactions of N243 with H152 and F153. **d)** Structural alignment of HTP-1 (grey) and HTP-1mut7 (orange) highlighting the movements observed at the end of β5.

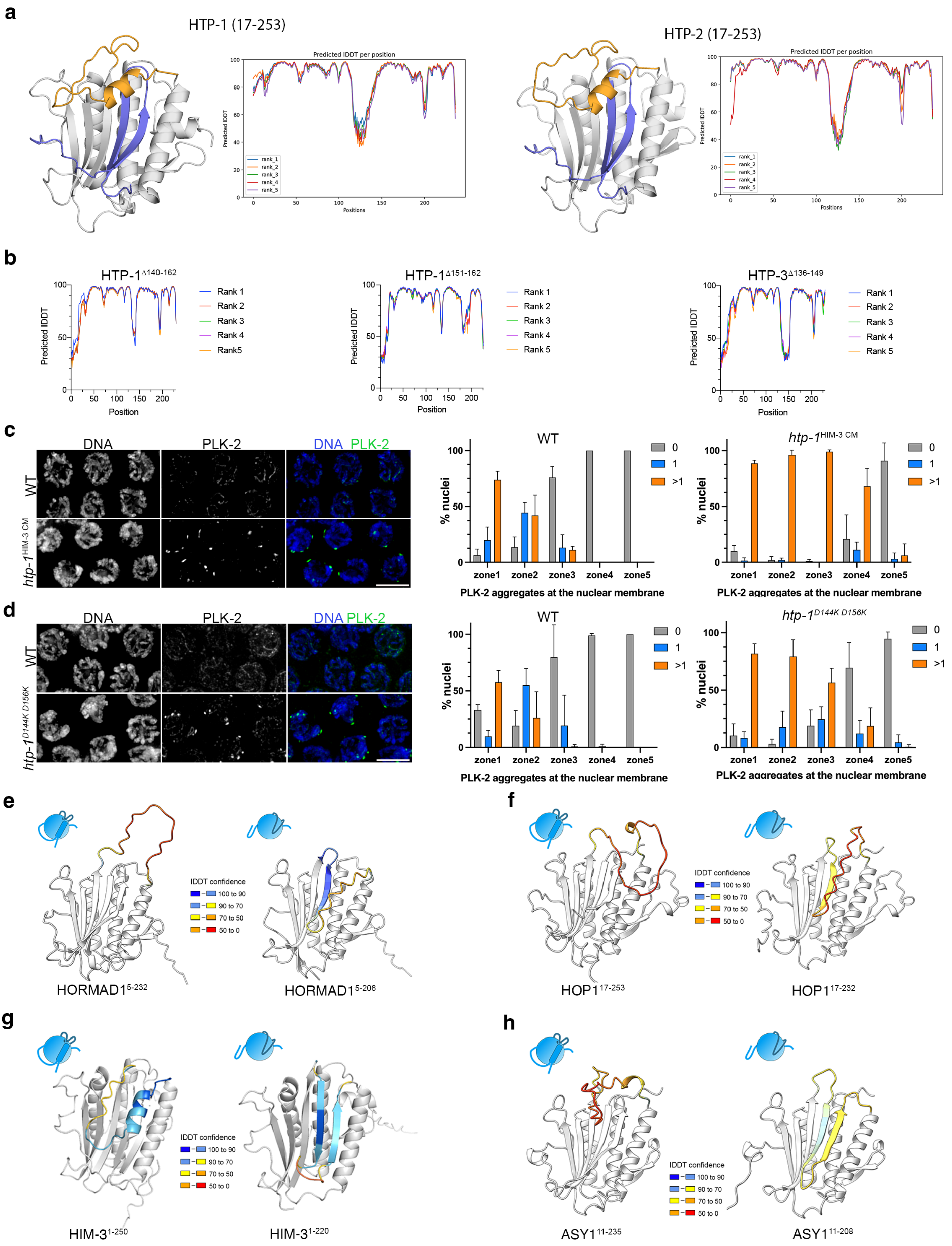
a
HTP-1 (17-253)
HTP-2 (17-253)
Predicted lDDT per position
Predicted lDDT
Positions
rank_1
rank_2
rank_3
rank_4
rank_5
b
HTP-1Δ140-162
HTP-1Δ151-162
HTP-3Δ136-149
Predicted lDDT
Position
Rank 1
Rank 2
Rank 3
Rank 4
Rank 5
c
DNA
PLK-2
DNA PLK-2
WT
htp-1HIM-3 CM
% nuclei
0
1
>1
zone1
zone2
zone3
zone4
zone5
PLK-2 aggregates at the nuclear membrane
d
htp-1D144K D156K
e
lDDT confidence
100 to 90
90 to 70
70 to 50
50 to 0
HORMAD15-232
HORMAD15-206
f
HOP117-253
HOP117-232
g
HIM-31-250
HIM-31-220
h
ASY111-235
ASY111-208

**Figure S4. a)** ColabFold predictions of HTP-1 and HTP-2 HORMA domains (residues 17-253) indicating the β5-αC loop in orange and the safety belt region in blue. Note that HTP-1 and HTP-2 HORMA domains are predicted to display an empty closed conformation. Graphs show the predicted lDDT per position for the five models (ranks 1-5), indicating high confidence in the position of the safety belt region. **b)** Graphs showing predicted lDDT per position for five models (ranks 1-5) of indicated HTP-1 and HTP-3 loop mutants (models shown in Fig 7b). **c-d)** Projections of pachytene nuclei of indicated genotypes stained with anti-PLK-2 antibodies and DAPI. Nuclei with more than 1 PLK-2 aggregate indicate high CHK-2 activity, 1 PLK-2 aggregate indicates intermediate CHK-2 activity and no PLK-2 aggregate indicates no CHK-2 activity. Graphs show quantification of the % of nuclei with a given number of PLK-2 aggregates in five zones along the germline. Both *htp-1*$^{HIM\text{-}3\ CM}$ and *htp-1*$^{D144K\ D156K}$ mutants accumulate nuclei with multiple PLK-2 aggregates. Number of nuclei (3 to 7 germlines per genotype) analysed per zone= 233, 207, 172, 163, 121 (WT); 183, 150,145, 98, 98 (*htp-1*$^{HIM\text{-}3\ CM}$); 176, 148, 129, 121, 96 (WT); 507, 412, 358, 296, 232 (*htp-1*$^{D144K\ D156K}$). **e-h)** ColabFold models of the HORMA domain of indicated mHORMADs indicating lDDT confidence values for the β5-αC loop. Model on the left corresponds to WT sequence and model on the right corresponds to structure lacking the safety belt. Note that in all cases deletion of the safety belt resulted in the interaction of the β5-αC loop with β5. Scale bar= 5 µm in all panels.
