## Supplemental Table 2 for "Two structurally mobile regions control the conformation and function of metamorphic meiotic HORMAD proteins"

| **Transgene** | **Genotype** |
| --- | --- |
| *fqSi6* | [*Phtp-1 htp-1 wild type 3’UTR htp-1*; *cb-unc-119(+)*] |
| *fqSi22* | [*Phtp-1 htp-1::6HIS 3’UTR htp-1*; *cb-unc-119(+)*] |
| *ieSi27* | [*Phtp-1 htp-2 N-terminus(1-41aa) htp-1(42-352aa) 3’ UTR htp-1; cb-unc-119(+)*] |
| *fqSi28* | [*Phtp-1 htp-1 (1-250aa) htp-2 C-terminus (251-352aa) 3’ UTR htp-1; cb-unc-119(+)*] |
| *fqSi30* | *[Phtp-1 htp-1 (1-41aa) htp-2 HORMA (42-250aa) htp-1(251-352aa) 3’ UTR htp-1*; *cb-unc-119(+)*] |
| *fqSi141* | [*Phtp-1 htp-1 (1-41aa) htp-2 HORMA A (42-90aa) htp-1 (91-352aa)::6HIS 3’UTR htp-1*; *cb-unc-119(+)*] |
| *fqSi181* | *[Phtp-1 htp-1(1-90aa) htp-2 HORMA B (91-165aa) htp-1 (166-352aa)::6HIS 3’UTR htp-1*; *cb-unc-119(+)*] |
| *fqSi167* | [*Phtp-1 htp-1 (1-165aa) htp-2 HORMA C (166-250aa) htp-1 (251-352aa)::6HIS 3’UTR htp-1*; *cb-unc-119(+)*] |

**Table S2. List of transgenes used in this study.**
