## Supplemental Table 1 for "Two structurally mobile regions control the conformation and function of metamorphic meiotic HORMAD proteins"

| **Strains** | **Genotype** | **Origin** |
| --- | --- | --- |
| N2  Bristol | Wild type | CGC |
| AV393 | *htp-1(gk174)/ nT1[unc-?(n754) let-? qIs50] (IV;V)* | (Martinez et al., 2005; Couteau et al.,2005) |
| AV276 | *syp-2(ok307) V/nT1[unc-?(n754) let-?(m435)] (IV;V)* | (Colaiácovo et al., 2003) |
| EG6699 | *ttTi5605 II MOSCI; unc119(ed3)III; oxEx1578* |  |
| ATG285 | *htp-1 (fq24 [htp-1::FLAG]) IV* | This study |
| ATG284 | *htp-2 (fq26 [htp-2::FLAG]) IV* | This study |
| ATG224 | *htp-2 (fq26[htp-2::FLAG])IV; htp-1(gk174)/ nT1 [unc-? (n754) let-? qls50] (IV;V)* | This study |
| ATG718 | *fqSi6 II; htp-1(gk174)* | This study |
| ATG711 | *fqSi27 II; htp-1(gk174)* | This study |
| ATG712 | *fqSi28 II; htp-1(gk174)* | This study |
| ATG713 | *fqSi30 II; htp-1(gk174)/ nT1 [unc-? (n754) let-? qls50] (IV;V)* | This study |
| ATG719 | *fqSi22 II; htp-1(gk174)* | (Ferrandiz et al., 2018) |
| ATG714 | *fqSi141 II; htp-1(gk174)* | This study |
| ATG715 | *fqSi181 II; htp-1(gk174)* | This study |
| ATG716 | *fqSi167 II; htp-1(gk174)/ nT1 [unc-? (n754) let-? qls50] (IV;V)* | This study |
| ATG540 | *htp-1 (syb1225 [htp-2]) IV/ tmC25 [unc-5(tmIs1241)] IV* | This study |
| ATG665 | *htp-1 (fq162 [htp-2::FLAG] IV/ tmC25 [unc-5(tmIs1241)] IV* | This study |
| ATG607 | *htp-1 (fq141 [htp-1Δ]) IV/ tmC25 [unc-5(tmIs1241)] IV* | This study |
| ATG669 | *htp-2 (fq163 [htp-2Δ]) IV* | This study |
| AV393 | *htp-1(gk174)/ nT1[unc-? (n754) let-? qIs50] (IV;V)* | Martinez-Perez et al 2005() |
| ATG717 | *htp-1 (syb4096 [E191A E200K]) IV* | This study |
| ATG643 | *htp-1 (fq152 [htp-1C239H N243H E245Q::FLAG]) IV/ tmC25 [unc-5(tmIs1241)] IV* | This study |
| *ATG650* | *htp-1 (fq154 [htp-1 M249KL250SC239HN243HE245Q::FLAG]) IV/ tmC25 [unc-5(tmIs1241)] IV* | This study |
| ATG656 | *htp-1 (fq157 [htp-1 D226G A230V C239H N243H E245Q M249K and L250S::FLAG])/ tmC25 [unc-5(tmIs1241)] IV* | This study |
| ATG677 | *htp-1 (fq24 [htp-1::FLAG]) IV; htp-2 (fq163 [Δhtp-2]) IV* | This study |
| ATG491 | *htp-1 (fq98 [htp-2 HORMA C E191-L250]) IV/ nT1[unc-? (n754) let-? qIs50] (IV;V)* | This study |
| ATG493 | *htp-1 (fq98 [htp-2 HORMA C E191-L250]) IV/ tmC25 [unc-5(tmIs1241)] IV* | This study |
| ATG565 | *htp-1 (fq128 [htp-2 HORMA C E191-L250::FLAG]) IV/ tmC25 [unc-5(tmIs1241)] IV* | This study |
| ATG588 | *htp-1 (fq98 [htp-2 HORMA C E191-L250]) IV; syp-2 (ok307) V/ nT1[unc-? (n754) let-? qIs50] (IV;V)* | This study |
| ATG582 | *htp-1 (syb1225 [htp-2]) IV; syp-2 (ok307)/ nT1[unc-? (n754) let-? qIs50] (IV;V)* | This study |
| AV402 | *htp-1(gk174) IV; syp-2(ok307) V)/ nT1[unc-? (n754) let-? qIs50] (IV;V)* | Martinez-Perez et al 2005() |
| AV276 | *syp-2 (ok307)/ nT1[unc-? (n754) let-? qIs50] (IV;V)* | This study |
| ATG542 | *htp-1 (fq115 [P151-E162]) IV/ tmC25 [unc-5(tmIs1241)] IV* | This study |
| ATG543 | *htp-1 (fq116 [R140-E162]) IV/ tmC25 [unc-5(tmIs1241)] IV* | This study |
| ATG670 | *htp-1 (fq164 [ΔP151-E162::FLAG]) IV/tmC25 [unc-5(tmIs1241)] IV* | This study |
| ATG671 | *htp-1 (fq165 [ΔR140-E162::FLAG]) IV/tmC25 [unc-5(tmIs1241)] IV* | This study |
| ATG691 | *htp-3 (fq175 [S136-V149 deletion])I/ hT2[bli-4(e937) let-?(q782) qIs48] (I;III)* | This study |
| ATG654 | *htp-1 (fq156 [htp-1^HIM-3CM^::FLAG]) IV/ tmC25 [unc-5(tmIs1241)]* | This study |
| ATG726 | *htp-3 (fq185 [htp-3 A137K]/ tmC18 [dpy-5(tmIs1236)] I* | This study |

**Table S1. List of strains used in this study.**
